# RBFOX2 restrains oncogenic cell plasticity in Group 3 medulloblastoma

**DOI:** 10.64898/2026.10.02.756284

**Authors:** Rajiv Kumar Sah, Xinmiao Yan, Lexane Louis, Naomi Gonzales, Uyanga Tsogt, Baoqi He, Lily Greer Hoefner, Loïc Lecomte, Mason Bartels, Charlotte A Le Roux, Anne C Starner, Irene Basili, Olivier Ayrault, Eric L. Van Nostrand, Olivier Saulnier, Jamie Zagozewski, Tamra E. Werbowetski-Ogilvie

## Abstract

Cancer cell plasticity underlies tumor progression, metastasis and therapeutic resistance, yet how RNA-binding proteins regulate malignant state transitions remains poorly defined. Here, we identify the splicing regulator RBFOX2 as a suppressor of oncogenic cell plasticity in Group 3 medulloblastoma (MB). OTX2 antagonizes RBFOX2 by directing an opposing alternative splicing program that mirrors developmental origins and controls cytoskeletal dynamics and cell state transitions. Mechanistically, OTX2 alters the RBFOX2 RNA-binding landscape, limiting RBFOX2 engagement at cytoskeletal loci and suppressing exon inclusion. RBFOX2 loss rewires splicing and transcriptional outputs to promote highly plastic proliferative, mesenchymal-like and undifferentiated cellular states associated with enhanced oncogenic properties. Consistent with these findings, RBFOX2^low^ states mark highly plastic cell populations in patient tumors and xenografts, positioning RBFOX2 as a determinant of malignant cell stability. Together, these findings establish RBFOX2 as a developmental gatekeeper that restrains malignant cell plasticity through coordinated control of cytoskeletal gene programs.

## Introduction

Brain tumors are the leading cause of pediatric cancer-related mortality with medulloblastoma (MB) representing the most common malignant primary brain tumor in children. Among the MB subgroups, Group 3 MBs are the most aggressive, accounting for approximately one quarter of cases and exhibiting the worst clinical outcomes, with infants displaying a 5-year overall survival rate of 45%^1^. Leptomeningeal dissemination along the brain and spinal cord is the primary cause of death in Group 3 MB patients^1,2^, yet fewer than 10% of MB studies address the mechanisms driving metastatic disease. Progress has been further limited by the scarcity of metastatic patient samples, resulting in substantial gaps in our understanding of the regulatory programs governing MB cell fate and tumor spread.

Group 3 MBs are thought to originate from cells in the developing human cerebellar rhombic lip (RL) that retain primitive oncogenic properties^3,4^. A defining feature of Group 3 MB is elevated or amplified expression of the key transcription factor OTX2. OTX2 sustains this primitive, stem-like state by repressing neurodevelopmental gene expression^4–7^ and by directing an oncogenic alternative splicing (AS) program in association with the Large Assembly of Splicing Regulators (LASR) complex^8^. These observations indicate that Group 3 MB maintenance depends on coordinated transcriptional and post-transcriptional regulation to preserve immature cellular identities. More broadly, they implicate RNA-binding proteins (RBPs) as key mediators of developmental splicing programs that control cell plasticity and phenotypic state transitions. As cellular plasticity is a central feature of tumor heterogeneity, metastasis and drug resistance^9^, defining how RBPs regulate AS programs to shape dynamic Group 3 MB cell fate transitions remains an important unanswered question.

The RNA-binding protein RBFOX2 controls diverse AS events that are critical for tumor progression^10–12^. As a member of the multimeric LASR complex^13^, RBFOX2 primarily functions as a splicing regulator but has also been linked to transcriptional regulation, RNA stability and broader RNA metabolic functions^14,15^. Notably, RBFOX2-dependent splicing programs have been implicated in metastatic progression and EMT-associated phenotypes across multiple tumor types^10,14,16–25^. RBFOX2 is essential for murine cerebellar development, where it regulates neuronal differentiation and lineage-specific splicing programs^26^; however, its roles in human cerebellar development and MB progression remain unexplored.

We previously demonstrated that OTX2 silencing decreases self-renewal and promotes neural differentiation^6,27^ reflecting a shift from a stem cell to a differentiated cell state. Notably, this transition is accompanied by increased RBFOX2 expression^8^, raising the possibility that RBFOX2 opposes OTX2-driven cell state programs. Given its established role as a critical splicing regulator, we hypothesized that RBFOX2 functions as a molecular anchor that constrains oncogenic cell plasticity by opposing OTX2-regulated AS events.

Here, we identify RBFOX2 as a developmental regulator of malignant cell-state stability in MB. We establish an antagonistic regulatory axis in which RBFOX2 counteracts OTX2 to control an AS program governing cytoskeletal organization. Opposing splicing activity signatures are also mirrored in human cerebellar developmental origins. We show that this interplay reshapes gene regulatory programs associated with proliferation, actin dynamics and mesenchymal-like transitions. Functionally, RBFOX2 suppresses proliferative, mesenchymal-like cellular states and metastatic competence while RBFOX2^low^ cells are enriched for undifferentiated, EMT-like and proliferative signatures in both preclinical animal models and patient tumors. Together, our findings identify RBFOX2 as a key regulator of malignant cell plasticity that operates through coordinated control of cytoskeletal gene splicing programs central to Group 3 MB progression.

## Results

### OTX2 and RBFOX2 co-regulate a cytoskeletal gene splicing program

Our previous work demonstrated that OTX2 interacts with members of the LASR complex and plays a key role in regulating Group 3 MB AS^8^. To define how OTX2 and LASR components cooperatively regulate AS, we silenced OTX2 and three core LASR members, HNRNPC, HNRNPM and RBFOX2, using siRNAs and performed RNA-sequencing (RNA-seq) in HDMB03 MB stem-cell enriched tumorspheres (Extended Data Figure 1a-d). This analysis identified 112 differentially spliced genes (DSGs) shared across all four perturbations, with over two-thirds of OTX2-regulated DSGs also regulated by at least one LASR component (Fig. 1a, Supplementary Table S1), including previously identified OTX2 targets such as *PPHLN1*, *MBD1*, *WNK1*, and *EPB41L3*^8^.

**Figure 1:**
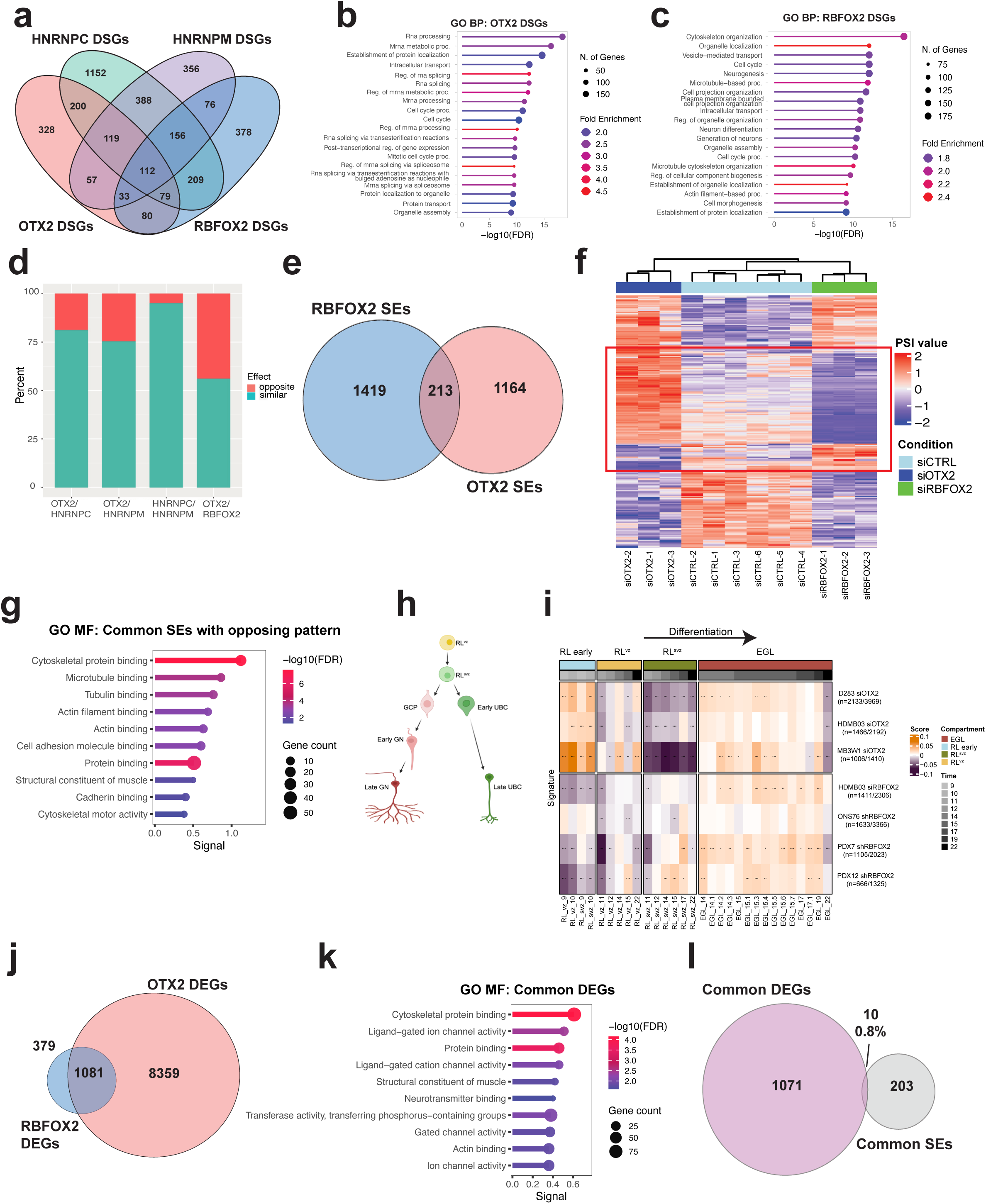
OTX2 and LASR proteins regulate common sets of splicing events. a. Venn diagram depicting overlap in OTX2- and LASR protein-regulated DSGs in HDMB03 tumorspheres, including 112 DSG shared between all datasets. Data represent n=3 biological replicates from each gene silencing condition. b-c. Lollipop plot depicting enrichment of GO Biological Processes (GO BP) associated with OTX2-regulated (b) and RBFOX2-regulated (c) DSGs. d. Bar graph depicting proportion of similar or opposing splicing patterns of shared splicing events regulated by OTX2 and LASR proteins. Similar splicing pattern indicates exon similarly excluded or included in both conditions. Opposing splicing pattern indicates exon excluded in one condition and included in the other condition. e. Venn diagram depicting 213 shared RBFOX2-regulated SE and OTX2-regulated SE in HDMB03 tumorspheres. SE=spliced exons. f. Heat map demonstrating the splicing patterns of the 213 shared RBFOX2-regulated and OTX2-regulated SE. Box highlights RBFOX2- and OTX2-regulated SE that have opposing splicing patterns. Data represent n=3 for siOTX2 and siRBFOX2 and n=6 for siCTRL treatments. PSI = percent spliced in, SE = spliced exons, CTRL = control, g. Lollipop plot demonstrating the GO Molecular Function (GO MF) enrichment for common OTX2- and RBFOX2-regulated SE with opposing splicing patterns. h. Schematic representation of human cerebellar lineages progressing from early to late developmental stages. Image created in BioRender. i. Heat map of OTX2- and RBFOX2-regulated splicing activity signatures across multiple MB cell lines and culture-adapted PDX MB cell lines projected onto developing human cerebellar data. Group 3 MB lines include D283, HDMB03, MB3W1 and culture-adapted PDX7. SHH MB lines include ONS76 and culture-adapted PDX12. Arrow denotes differentiation. *** FDR < 0.001, ** FDR < 0.01, and * FDR < 0.05. RL = rhombic lip, RL^VZ^ = rhombic lip ventricular zone, RL^SVZ^ = rhombic lip subventricular zone, EGL = external granule layer. j. Venn diagram depicts the overlap between OTX2- and RBFOX2-regulated DEGs in HDMB03 tumorspheres. DEG = differentially expressed genes. k. Lollipop plot demonstrating the GO Molecular Function (GO MF) enrichment for common shared OTX2- and RBFOX2-regulated DEG. l. Venn diagram depicting the overlap between common OTX2- and RBFOX2 regulated DEGs and common OTX2- and RBFOX2-regulated SEs in HDMB03 tumorspheres.

Globally, OTX2-regulated DSGs were enriched for RNA processing and splicing functions, whereas HNRNPC- and HNRNPM-regulated events exhibit highly similar splicing signatures enriched for cell cycle-associated genes (Fig. 1b, Extended Data Fig. 1e-f). In contrast, RBFOX2 controls a distinct AS signature strongly enriched for cytoskeletal reorganization (Fig. 1c). Directional analysis of shared splicing events (exon inclusion vs. skipping) revealed high concordance between OTX2, HNRNPC and HNRNPM (Fig. 1d). Notably, nearly half of exons co-regulated by OTX2 and RBFOX2 exhibited opposing splicing outcomes indicating a unique antagonistic regulatory relationship (Fig. 1d).

Specifically, OTX2 and RBFOX2 co-regulated a shared set of 213 alternatively spliced exons (SEs) (Fig. 1e, Supplementary Table S2). Of these, 99 events displayed opposing splicing outcomes where, where OTX2 loss primarily promotes exon inclusion (higher percent spliced in (PSI)), and RBFOX2 loss promotes exon skipping or the shorter variant (lower PSI) (Fig. 1f). These antagonistically regulated SEs were significantly enriched for genes that control cytoskeletal organization and cell adhesion, with many having known functional roles in metastasis (Fig. 1g). Exons regulated in the same direction showed minimal enrichment for cytoskeletal functions (Extended Data 1g-h).

To assess the broader relevance of this antagonistic splicing program in primitive stem cell populations, we extended our analyses to five additional OTX2- or RBFOX2-silenced MB models, including *in vitro* cell cultures and culture-adapted patient-derived xenograft (PDX) lines spanning multiple MB subgroups. Projection of OTX2- and RBFOX2-regulated splicing activity signatures onto a human developmental cerebellar dataset^28^ revealed that OTX2-associated splicing activity aligned with the primitive RL stem cell populations (RL early) at 9-10 weeks post conception, whereas RBFOX2-dependent splicing activity signatures were actively suppressed in this compartment (Fig. 1h-i). These findings indicate that the functional OTX2-RBFOX2 antagonism reflects conserved regulatory programs recapitulated during early human cerebellar development.

Finally, comparison of differentially expressed genes (DEGs) across all perturbations revealed minimal overlap between gene expression changes and AS regulation (Extended Data Fig. 2a-l, Supplementary Table S3). OTX2 and RBFOX2 shared 1081 DEGs (Fig. 1j) that are also highly enriched for cytoskeletal dynamics (Fig.1k). However, only 0.8% of the 213 shared OTX2- and RBFOX2-regulated SEs were also differentially expressed (Fig. 1l) demonstrating that our antagonistic cytoskeletal gene program is primarily driven by AS rather than changes in transcript abundance.

Collectively, these results identify a distinct antagonistic AS relationship between OTX2 and RBFOX2, centered on cytoskeletal-associated genes and mirrored in MB developmental origins, suggesting that RBFOX2-mediated AS has putative suppressive functions in MB stem cells.

### OTX2 modulates RBFOX2 RNA binding to specify cytoskeletal gene splicing outcomes

We previously demonstrated that OTX2 silencing decreases stemness and promotes neural differentiation, a transition accompanied by increased RBFOX2 levels as well as changes in localization^6,8,27^. Given the antagonistic splicing patterns observed between OTX2 and RBFOX2, we hypothesized that OTX2 modulates RBFOX2 engagement with target RNAs. To test this, we performed RNA-binding protein motif enrichment analysis following OTX2 and RBFOX2 silencing in Group 3 MB tumorspheres. The canonical RBFOX2 binding motif was most significantly enriched downstream of exons excluded in siRNA control (siCTRL) cells relative to siRNA silenced OTX2 (siOTX2) cells (Extended Data 3a). In contrast, this motif was enriched downstream of exons included in siCTRL relative to siRNA silenced RBFOX2 (siRBFOX2) cells (Extended Data 3b), consistent with established RBFOX2 positional effects on exon inclusion and skipping events^14^ (Fig. 2a).

**Figure 2:**
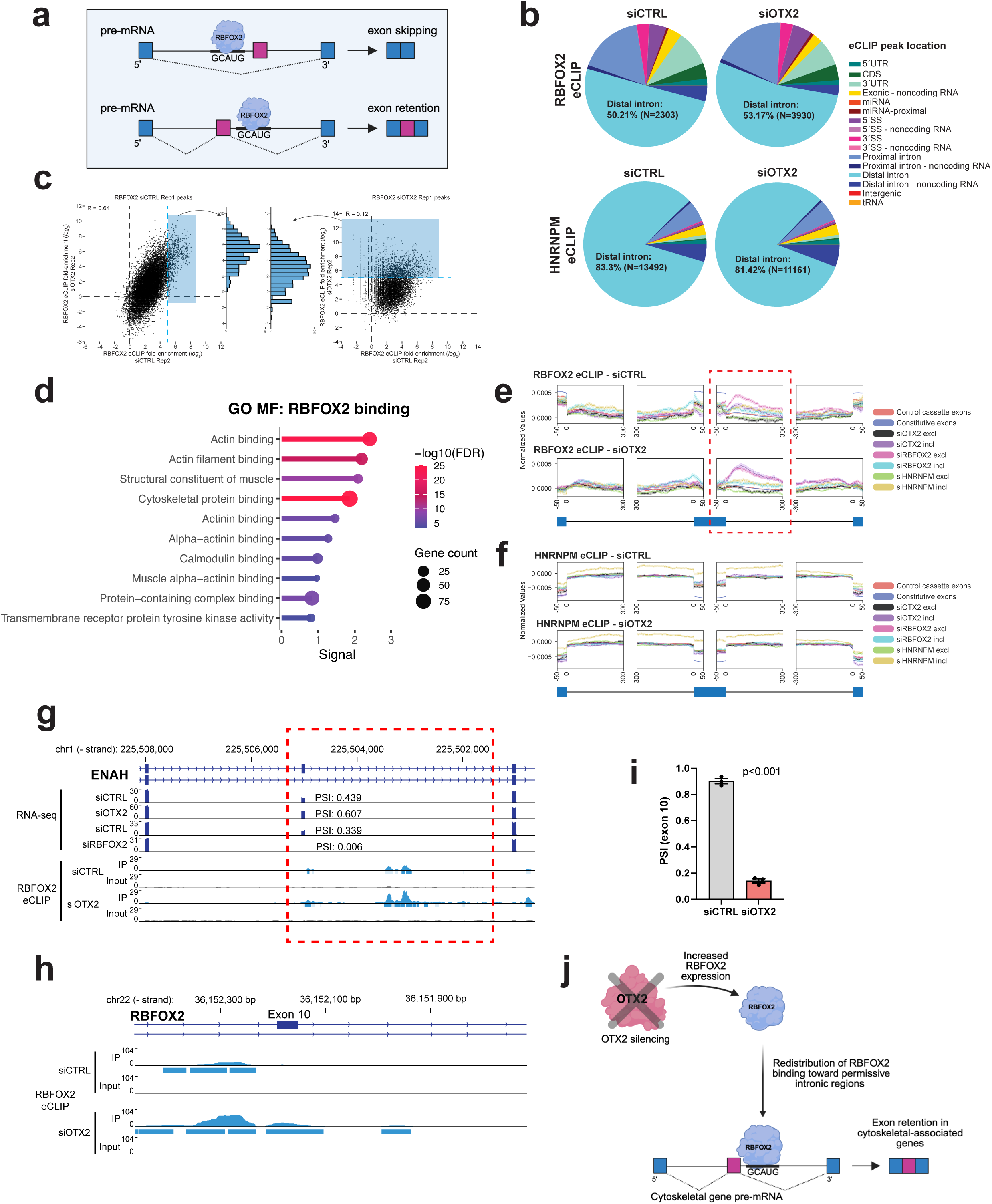
OTX2 reshapes the RBFOX2 binding landscape to control cytoskeletal splicing. a. Schematic figure depicting binding of RBFOX2 to consensus motifs upstream of target exon promotes exon skipping, while RBFOX2 binding to consensus motifs downstream promote exon retention. Image created in BioRender. b. Pie charts indicate the fraction of reproducible peaks from RBFOX2 or HNRNPM eCLIP that overlap indicated gene annotation regions. c. For each peak identified in one replicate of RBFOX2 eCLIP performed in siCTRL or siOTX2 conditions, points indicate the fold-enrichment in IP versus input for (x-axis) Replicate 2 in siCTRL or (y-axis) Replicate 2 in siOTX2. Histograms indicate the distribution of fold-enrichments observed in the opposite condition (e.g. siOTX2 enrichment for siCTRL peaks or siCTRL enrichment for siOTX2 peaks). d. Enrichment of GO Molecular Functions (GO MF) for genes associated with increased RBFOX2 peaks following OTX2 silencing in HDMB03 tumorspheres. e-f. Splicing maps indicate enrichment of eCLIP binding for (e) RBFOX2 and (f) HNRNPM for exons with altered splicing upon knockdown of OTX2, RBFOX2, or HNRNPM, along with control constitutive and alternative exons. Regions shown include 300 nt of proximal intron and 50 nt of flanking exonic region for the alternative exon as well as the upstream and downstream exon. Box highlights regions of RBFOX2 enrichment downstream of OTX2 included exons. g. Genome browser view of ENAH SE. RNA-seq tracks (boxed region) show increased exon inclusion following OTX2 depletion (siOTX2; increased PSI) and exon skipping following RBFOX2 depletion (siRBFOX2; decreased PSI) Corresponding eCLIP tracks (boxed region) demonstrate increased RBFOX2 binding downstream of ENAH SE in siOTX2 cells relative to siCTRL. PSI values represent the mean of three biological RNA-seq replicates; displayed tracks are from a representative replicate. PSI = percent spliced in, SE = spliced exon. h. Genome browser figure demonstrating enrichment of RBFOX2 upstream of *RBFOX2* exon 10 upon OTX2 silencing in HDMB03 tumorspheres. i. Quantification of PSI value from HDMB03 RNA-sequencing data showing increased exclusion of RBFOX2 exon 10 in OTX2-silenced cells. n=3 biological replicates. P<0.001. PSI = percent spliced in. j. Schematic illustrating the proposed mechanism by which OTX2 regulates RBFOX2-dependent alternative splicing. OTX2 silencing in Group 3 MB tumorspheres increases RBFOX2 expression and promotes enrichment downstream of cytoskeletal-associated target exons, increasing exon retention in accordance with canonical RBFOX2 binding patterns. Image created in BioRender.

To determine whether OTX2 regulates AS through direct RNA binding, we performed enhanced crosslinking and immunoprecipitation (eCLIP) of OTX2 in HDMB03 tumorspheres. Immunoblot validation indicated successful immunoprecipitation of OTX2 (Extended Data 3c), but we observed relatively low RNA yield. Standard eCLIP peak-calling criteria identified only 16 significant peaks (Extended Data 3d) many of which localized at rRNA pseudogenes or other common artifact regions. This signal was comparable to previous observations in IgG controls and notably less than observed from bona fide RBPs^29^ and V5 or FLAG antibody pulldowns in wild-type cells^30^. These findings argue against widespread direct interaction between OTX2 and differentially spliced exons. Analysis of mapped reads did not indicate substantial enrichment for distinct RNA classes^31^ but we did observe a moderate (2.0 to 2.7-fold) increase in reads at intergenic regions or loci antisense to annotated genes (Extended Data 3e), possibly reflecting transient co-transcriptional interactions due to OTX2 association with chromatin.

As these data suggest that OTX2 does not directly bind to RNA to regulate AS events, we next tested whether OTX2 influences RBFOX2 binding to target transcripts by performing eCLIP for RBFOX2 in HDMB03 tumorspheres following OTX2 silencing, with HNRNPM evaluated for comparison. Consistent with ENCODE datasets, most RBFOX2 and HNRNPM peaks localized predominantly to distal intronic regions (Fig. 2b). Notably, OTX2 knockdown largely preserves or increases the enrichment at RBFOX2 peaks that is observed in control cells (Fig. 2c, left) while also revealing novel RBFOX2 binding peaks (Fig. 2c, right). Genes exhibiting increased RBFOX2 binding upon OTX2 silencing were significantly enriched for functions related to cytoskeletal dynamics, with a particular emphasis on actin remodeling (Fig. 2d, Supplementary Table S4).

To assess binding preferences globally, we generated splicing maps for RBFOX2 and HNRNPM (Fig. 2e-f). OTX2 silencing increases RBFOX2 binding downstream of cytoskeletal transcripts promoting exon retention, (Fig. 2e), including the known RBFOX2 target ENAH Actin Regulator (*ENAH*)^17^ (Fig. 2g). This binding pattern was not observed for HNRNPM (Fig. 2f), demonstrating that OTX2 modulates RBFOX2-RNA engagement of select cytoskeletal targets. In addition, OTX2 loss increased RBFOX2 binding to its own transcript at alternative exon 10 (Fig. 2h) suggesting activation of an autoregulatory loop controlling RBFOX2 isoform usage. In support of these findings, OTX2 silencing promotes *RBFOX2* isoform switching toward exon 10 skipping (reduction in PSI value) in Group 3 MB tumorspheres (Fig. 2i). Exon 10 skipping is known to generate a cytoplasmic RBFOX2 isoform^10,32^, and this is consistent with our previous findings demonstrating a RBFOX2 subcellular localization shift in OTX2-silenced cells^8^.

Together, these findings demonstrate that OTX2 alters the RBFOX2 RNA-binding landscape to direct opposing AS outcomes (Fig. 2j). This OTX2-dependent tuning of RBFOX2 binding provides the mechanistic basis for the observed OTX2-RBFOX2 antagonistic control of cytoskeletal gene splicing programs in stem-cell enriched Group 3 MB cell populations.

### RBFOX2 suppresses oncogenic properties of Group 3 medulloblastoma *in vitro*

Our RBFOX2-mediated splicing activity signature is conserved across MB models, enriched for genes associated with actin remodeling, and downregulated in primitive cerebellar stem cell populations. Based on these findings, we hypothesize that RBFOX2 functions to restrain oncogenic properties in Group 3 MB stem cells. To test this, we performed reciprocal gain and loss of function studies in HDMB03 Group 3 MB cells. RBFOX2 was silenced using siRNAs (Fig. 3a) in both tumorspheres and monolayer cultures in stem cell-enriched conditions reflecting differences in cell adhesion properties.

**Figure 3:**
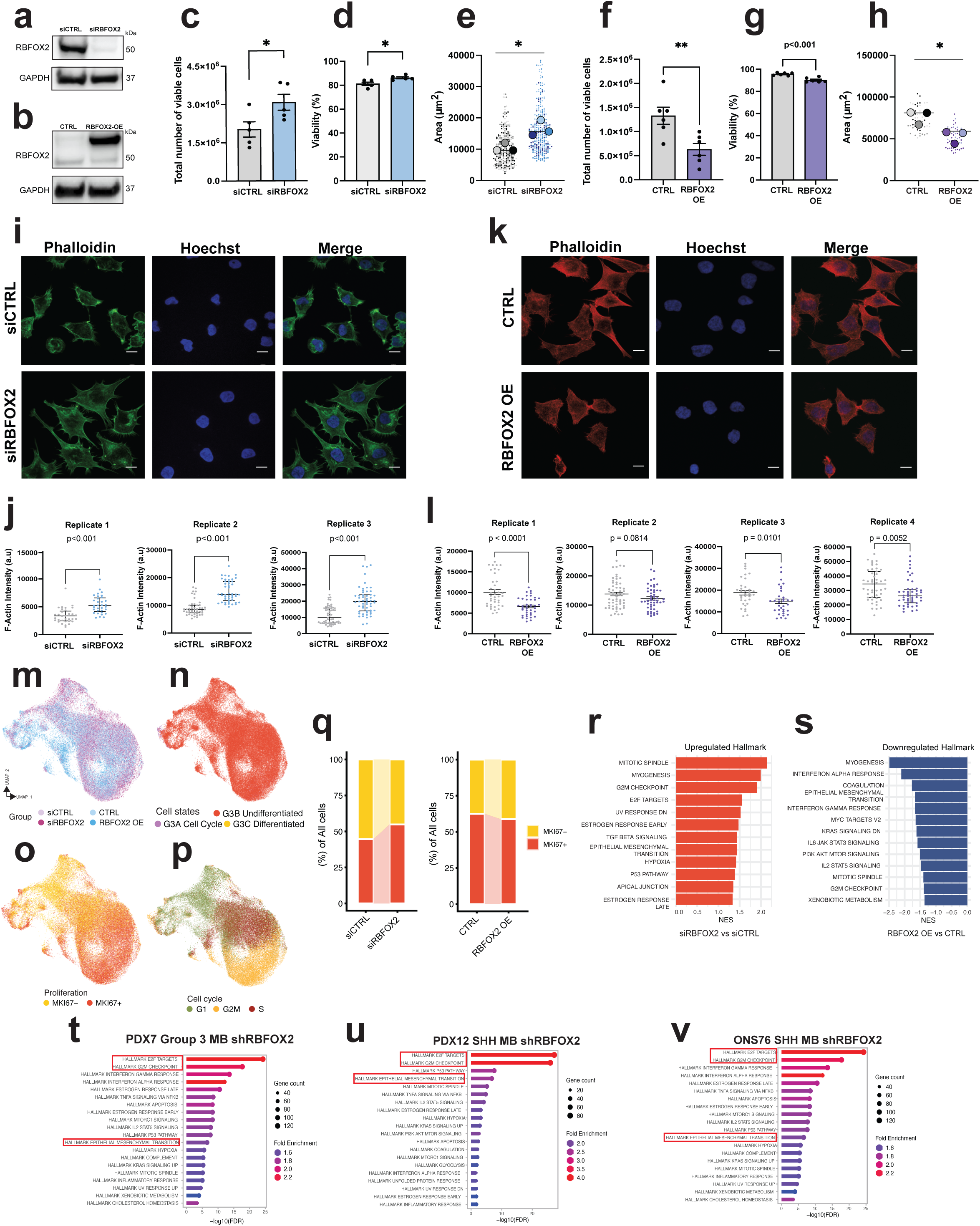
RBFOX2 inhibits oncogenic properties in Group 3 MB cells. a-b. Immunoblot depicting RBFOX2 silencing (a) and RBFOX2 overexpression (b) in HDMB03 cells. GAPDH serves as a loading control. c-e. Quantification of cell number (c), viability (d), and tumorsphere size (e) in siCTRL and siRBFOX2 treated HDMB03 tumorspheres. Error bars (c-d) represent s.e.m. (n=5 biological replicates). Statistical significance was determined using an unpaired t-test. Cell number: *P=0.040, viability: *P=0.0128. For tumorsphere size analysis (e), small individual dots represent single tumorspheres (technical replicates) nested within three independent experiments (indicated by three distinct colors). Larger circles show the mean tumorsphere area per replicate experiment, horizontal lines indicate mean ± SEM, and statistical significance was determined using a nested t-test. *P=0.0208. f-h. Quantification of cell number (f), viability (g), and tumorsphere size (h) following RBFOX2 overexpression (RBFOX2 OE) in HDMB03 tumorspheres. Error bars (f-g) represent s.e.m (n=6 biological replicates). Statistical significance determined using an unpaired t-test. Cell number: **P=0.0091, viability: P<0.001. For tumorsphere size analysis (h), small individual dots represent single tumorspheres (technical replicates) nested within three independent experiments (indicated by three distinct colors). Larger circles show the mean tumorsphere area per replicate experiment, horizontal lines indicate mean ± SEM, and statistical significance was determined using a nested t-test. *P=0.0229. i. Representative immunofluorescent images depicting phalloidin staining on siCTRL and siRBFOX2 treated HDMB03 cells. Hoechst staining indicates nuclei. Green channel: phalloidin, blue: nuclei. j. Quantification of F-actin intensity following phalloidin staining in siCTRL and siRBFOX2 treated HDMB03 cells. Results from three independent replicates shown. Error bars represent median with interquartile range. Statistical significance was determined using a Mann-Whitney test. Replicate 1: P <0.001, Replicate 2: P <0.001. Replicate 3: P <0.001. k. Representative immunofluorescent images depicting phalloidin staining on CTRL and RBFOX2 OE treated HBMB03 cells. Hoechst staining indicates nuclei. Red channel: phalloidin, blue: nuclei. l. Quantification of F-actin intensity following phalloidin staining in CTRL and RBFOX2 OE treated HDMB03 cells. Results from four independent replicates shown. Error bars represent median with interquartile range. Statistical significance was determined using a Mann-Whitney test. Replicate 1: P <0.001, Replicate 2: P = 0.0814. Replicate 3: P = 0.0101. Replicate 4: P= 0.0052. m-p. UMAP of the scRNA-seq datasets (including siRBFOX2, siCTRL, RBFOX2 OE, and CTRL) showing single-cell profiles colored by sample (m), cell state (n), proliferation status (o), and cell-cycle phase (p). q. Composition of proliferation cell states between siCTRL and siRBFOX2 (left), and between CTRL and RBFOX2 OE (right). Colors correspond to the proliferation states shown in (o). r-s. Significantly enriched upregulated Hallmark pathways in siRBFOX2 versus siCTRL cells (r) and significantly enriched downregulated Hallmark pathways in RBFOX2 OE versus CTRL cells (s). t-v. Lollipop plots depicting Hallmark enrichment for RBFOX2-regulated DEGs in culture-adapted PDX7 Group 3 MB (t), culture adapted PDX12 SHH MB (u), and ONS76 SHH MB (v).

In parallel, RBFOX2 was overexpressed (RBFOX2 OE) using lentiviral delivery of full-length RBFOX2 cDNA. Successful RBFOX2 overexpression was also confirmed by immunoblot (Fig. 3b).

RBFOX2 silencing significantly increased total cell number and viability in both culture conditions (Fig. 3c-d, Extended Data 4a-b). Consistent with these findings, RBFOX2-silenced cells generated larger tumorspheres (Fig. 3e). Conversely, RBFOX2 OE produced the opposite effect, significantly reducing total cell number, viability and tumorsphere size (Fig. 3f-h).

Given the strong cytoskeletal/actin remodeling signatures observed in RBFOX2 silenced cells, we next examined RBFOX2 regulation of actin organization by phalloidin staining. RBFOX2 silenced cells exhibited marked cytoskeletal reorganization including increased cell spreading, pronounced elongation, and more irregular, protrusive borders (Fig. 3i). Quantitative analysis confirmed a significant increase in phalloidin intensity, reflecting redistribution of F-actin, increased cell area, and reduced circularity in siRBFOX2 cells, suggesting a shift toward a more mesenchymal-like contractile phenotype (Fig. 3j, Extended Data 4c). In contrast, RBFOX2 OE resulted in decreased F-actin intensity, reflecting cytoskeleton disassembly (Fig. 3k-l), without significant changes in cell area or morphology (Extended Data 4d).

Finally, to evaluate the impact of RBFOX2 on malignant cell states, we performed single cell RNA-sequencing (scRNA-seq) on tumorspheres derived from both RBFOX2-silenced and overexpressing cells (Fig. 3m-p, Extended Data 4e-i). Individual cells were assigned to Group 3 MB transcriptional metaprograms defined by Hovestadt *et al*^33^ representing cell state or cycling status. As expected, the undifferentiated program was dominant under stem cell-enriched culture conditions (Fig. 3n) with primitive cells spanning multiple cell cycle phases (Fig. 3o-p). Consistent with functional assays, RBFOX2 silencing increased the proportion of Ki67+ proliferating cells and G2M-phase cells while decreasing cells in G1, indicating a shift towards a more proliferative cellular state (Fig. 3q, Extended Data 4e-g). This pattern was reversed when RBFOX2 was overexpressed (Fig. 3q, Extended Data 4e-g). Hallmark pathways associated with proliferation and mesenchymal transition were reciprocally regulated by RBFOX2 gain and loss of function (Fig. 3r-s, Supplementary Table S5), suggesting that RBFOX2 influences coordinated transcriptional programs linked to MB stem cell plasticity^9,34,35^. Notably, the myogenesis gene set, which in this context globally reflects contractile actin remodeling and cytoskeletal reorganization programs, exhibited one of the strongest reciprocal responses, showing robust activation upon RBFOX2 silencing and suppression following overexpression (Fig. 3r-s, Extended Data 4h-i, Supplementary Table S5-7).

Having functionally assessed the role of RBFOX2 in Group 3 MB, we examined whether RBFOX2’s regulatory influence extends across additional Group 3 MB models and MB subgroups. These models include the culture-adapted PDX Group 3 MB line, PDX7, SHH MB ONS76 cells and the culture-adapted SHH PDX MB line, PDX12. Across all models, RBFOX2 loss consistently enriched for hallmark EMT and cell cycle signatures, mirroring cell-state shifts observed in our single nuclei dataset and functional assessments (Fig. 3t-v). Collectively, these *in vitro* data demonstrate that RBFOX2 restrains oncogenic cell-state transitions in MB, limiting the emergence of highly proliferative and EMT-like cellular states consistent with increased malignant plasticity.

### RBFOX2 limits malignant cell plasticity and metastatic competence in MB

Increasing evidence links metastatic competence to highly plastic tumor cell states enriched for stem-like, proliferative and hybrid-EMT programs^9,34,35^. Because RBFOX2 constrained the acquisition of these phenotypes *in vitro*, we investigated whether RBFOX2 regulates cellular plasticity and Group 3 MB leptomeningeal spread *in vivo*. First, we stably knocked out RBFOX2 using CRISPR-Cas9 and two sgRNAs. Consistent with transient silencing (Fig. 3), stable RBFOX2 depletion significantly enhanced cell number and viability *in vitro* (Extended Data 5a-d). However, this depletion did not further increase the highly aggressive tumor growth or leptomeningeal spread *in vivo* (Extended Data 5e-f), suggesting that RBFOX2 loss is sufficient to establish a highly, plastic state, but cannot further potentiate malignant progression.

To determine whether sustained RBFOX2 expression suppresses tumor progression *in vivo,* we injected either 1.5X10^4^ or 1.5X10^3^ stable HDMB03 RBFOX2 OE LV-CMV5-Luc2-GFP cells into the cerebellums of NSG mice and tracked tumor growth and metastasis using bioluminescent imaging (BLI). In contrast to RBFOX2 depletion, RBFOX2 OE significantly inhibited tumor progression. At day 15, primary tumors were evident in control (CTRL) and RBFOX2 OE mice, but metastatic dissemination was observed exclusively in CTRL animals (Fig. 4a). RBFOX2 OE tumors were also significantly smaller than CTRL tumors (Fig. 4b, Extended Data 5g-h). This translated to a significant extension in survival of RBFOX2 OE animals compared to CTRLs (Fig. 4c). Further evaluation of metastatic growth was conducted on tumors collected at end point using mitochondria staining for human cells (Fig. 4d). Only one small metastatic lesion was observed in a single RBFOX2 OE sample while metastatic lesions in the CTRL group were larger and more dispersed throughout the spinal leptomeninges (Fig. 4d).

**Figure 4:**
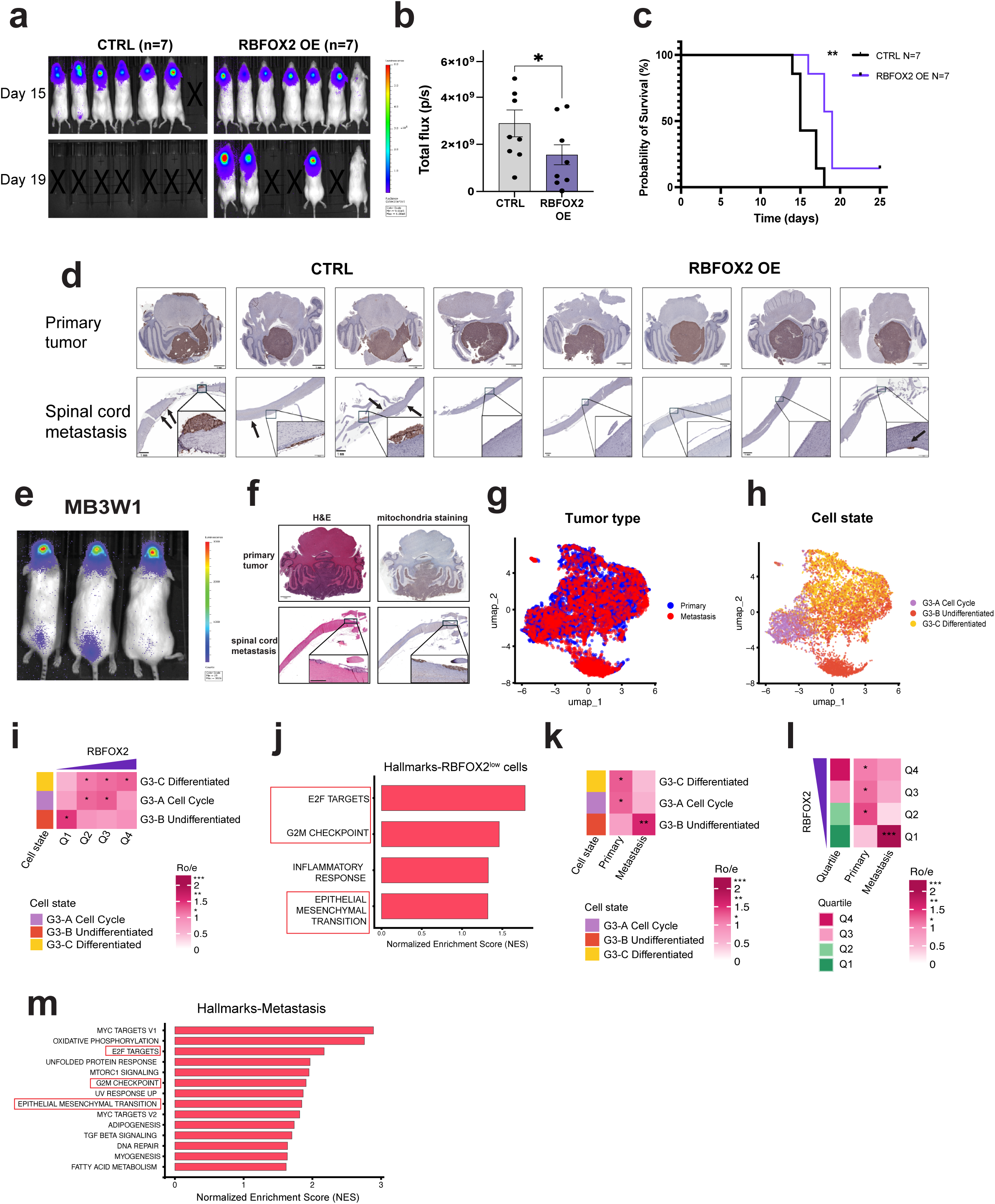
Sustained RBFOX2 limits malignant cell plasticity and metastatic competence *in vivo*. a. Bioluminescence imaging of NSG mice following intracerebellar injection of 1.5X10^4^ CTRL or RBFOX2 OE LV-CMV5-Luc2-GFP HDMB03 cells. Imaging was performed at day 15 and day 19 to visualize and track tumor growth. b. Quantification of tumor volume in animals injected with 1.5X10^4^ CTRL and RBFOX2-OE cells, calculated by BLI total flux (p/s). Statistical significance was determined by a Mann-Whitney test. n=8 or n=9 per group, *P=0.0464. c. Kaplan–Meier curves following transplantation of NSG mice with CTRL HDMB03 cells (n=7) and RBFOX2 OE HDMB03 cells (n=7). Statistical significance was determined by a log-rank test. **P = 0.0045. d. Mitochondrial staining on CTRL or RBFOX2 OE samples, highlighting the increased presence and volume of metastatic tumors in CTRL animals compared to RBFOX2 OE. Areas of metastatic tumor growth indicated with arrows. Tissue collected at survival endpoint for both groups. e. Representative bioluminescent imaging of primary and metastatic tumors arising from MB3W1 LV-CMV5-Luc2-GFP Group 3 MB cells injected into the cerebellum of NSG mice. f. Representative H&E staining (left) and mitochondrial IHC (right) showing primary and metastatic tumor growth from MB3W1 xenografts. Scale bars: 1 mm. g-h. UMAPs showing the snRNA-seq profiles of MB3W1 MB xenografts, with cells colored by tumor cell type (g) and cell states (h) annotated based on Hovestadt et al. derived metaprograms. G3=Group 3 MB. i. Cell state composition across RBFOX2 quartiles from lowest Q1 to highest Q4. ***Ro/e >3, ** Ro/e 1.5-2. *Ro/e >1. G3=Group 3 MB. j. Hallmark GSEA comparing RBFOX2^low^ (Q1) versus RBFOX2^high^ (Q4) cells in both primary and metastatic tumors. Boxes highlight EMT and cell cycle hallmarks. k. Cell state composition in primary versus metastatic tumor compartments. ***Ro/e >3, ** Ro/e 1.5-2. *Ro/e >1. G3= Group 3 MB. l. Quartile RBFOX2 cell composition between primary and metastatic MB. ***Ro/e >3, ** Ro/e 1.5-2. *Ro/e >1. m. Hallmark GSEA comparing metastatic versus primary tumor cells. Boxes highlight EMT and cell cycle hallmarks.

Next, we assessed the relationship between endogenous RBFOX2, stemness and metastasis in Group 3 MB tumors. Due to the paucity of metastatic patient samples, we performed single nuclei RNA-sequencing (snRNA-seq) on primary and microdissected metastatic Group 3 MB tumor cells derived from MB3W1 GFP-luciferase xenografts. This highly aggressive model has been shown to faithfully recapitulate the features of Group 3 MB patient tumors (Fig. 4e-f)^5,36,37^. In our dataset, clusters were also defined based on Group 3 MB metaprograms previously annotated by Hovestadt *et al*^33^ reflecting cell state (Fig. 4g-h). We first stratified all cells based on a continuum of low to high RBFOX2 expression. RBFOX2^low^ cells are significantly enriched for the undifferentiated state, as well as hallmark EMT and cell cycle signatures (Fig. 4i-j, Supplementary Table S8). Next, we examined the proportions of specific cell states in metastatic vs primary cells. Strikingly, RBFOX2^low^ cells are markedly overrepresented in the undifferentiated and metastatic cell population compared to primary tumor cells (Fig. 4k-l). Furthermore, the RBFOX2^low^ metastatic tumor cells are significantly enriched for EMT- and cell cycle-associated hallmark pathways, consistent with our gain and loss of function data *in vitro* (Fig. 4m, Supplementary Table S9). The co-enrichment of EMT-associated and proliferative programs suggests that RBFOX2^low^ metastatic cells occupy a highly plastic cellular state^9,34,35^.

Collectively, these *in vivo* results position RBFOX2 as a previously unrecognized gatekeeper of malignant cell state stability, rather than a linear driver of tumor progression, in Group 3 MB, restricting the emergence of highly plastic tumor cell populations enriched for EMT-like programs, undifferentiated cell states and increased metastatic competence.

### A conserved RBFOX2-regulated cytoskeletal splicing module is maintained across MB subgroups

To identify functionally relevant RBFOX2-regulated splicing events, we focused on transcripts exhibiting robust exon skipping following RBFOX2 silencing. We selected seven candidate cytoskeletal-associated genes including *ENAH*, LIM And Calponin Homology Domains 1 (*LIMCH1*), myosin light chain 6 (*MYL6*), abelson interactor 1 (*ABI1*), myosin phosphatase Rho-interacting protein (*MPRIP*), calsyntenin 1 (*CLSTN1*) and cortactin (*CTTN*). These genes have established roles in cytoskeletal dynamics, cell motility and have been identified as RBFOX2 targets in other malignancies^10,21,25,38–40^. RNA-seq (Fig. 5a) and RT-PCR validations (Fig. 5b, Extended Data 6a) revealed significant reduction in PSI, indicating exon exclusion following RBFOX2 silencing in both HDMB03 and MB3W1 stem cell-enriched tumorspheres. Consistent with the antagonistic OTX2-RBFOX2 regulatory axis, OTX2 loss predominantly promoted exon inclusion at these same target gene loci (Extended Data Fig. 6b).

**Figure 5:**
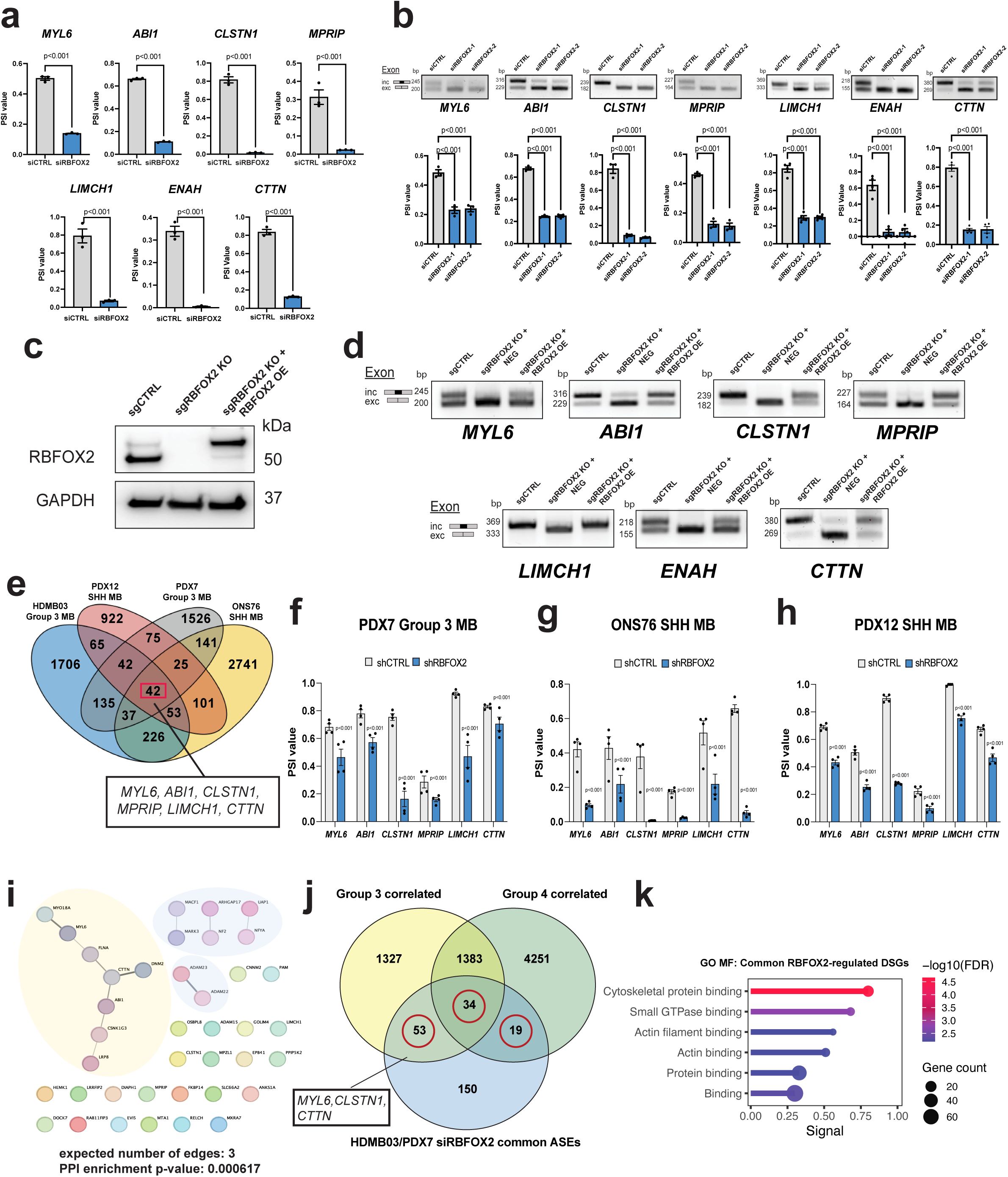
RBFOX2 regulates a conserved cytoskeletal splicing module across MB subgroups. a. Quantification of splicing alterations of select cytoskeletal remodeling genes from RNA-sequencing following RBFOX2 silencing in HDMB03 tumorspheres. Error bars represent s.e.m. (n=3 biological replicates). Values P<0.001 for all genes examined. Statistical significance was determined using FDR-adjusted p-values from RNA-sequencing data. PSI = percent spliced in. b. RT-PCR validation (upper) and associated quantification (lower) of alternatively spliced exon levels for select cytoskeletal remodeling genes in RBFOX2-silenced HDMB03 tumorspheres. Evaluation carried out with 2 independent RBFOX2 siRNAs. Error bars represent s.e.m. (n=3 biological replicates). Statistical significance determined using one-way ANOVA followed by a Dunnett’s test for multiple comparisons. P<0.001 for all genes examined. PSI = percent spliced in. c. Immunoblot depicting loss of RBFOX2 expression following CRISPR KO in HDMB03 cells and RBFOX2 re-expression in the KO background relative to control HDMB03 cells. GAPDH serves as a loading control. d. RT-PCR showing cytoskeletal gene exon exclusion (short isoform) is favored in RBFOX2 KO cells (sgRBFOX2 KO + NEG). Exon retention (long isoform) is rescued in RBFOX2 KO cells re-expressing RBFOX2 (sgRBFOX2 KO + RBFOX2 OE) demonstrating RBFOX2 splicing specificity. NEG = negative control. e. Venn diagram showing 42 shared RBFOX2-regulated spliced exons, including *MYL6*, *ABI1*, *CLSTN1*, *MPRIP*, *LIMCH1*, and *CTTN* between Group 3 MB models (HDMB03 and culture-adapted PDX7) and SHH MB models (ONS76 and culture-adapted PDX12). f-h. Quantification of splicing alterations of select cytoskeletal remodeling genes from RNA-sequencing following RBFOX2 silencing in the Group 3 MB culture-adapted PDX model, PDX7 (f) the SHH MB cell line, ONS76 (g) and the SHH MB culture-adapted PDX model, PDX12 (h). Error bars represent s.e.m. n=4 biological replicates for each. P<0.001 for all genes examined. Statistical significance was determined using FDR-adjusted p-values from RNA-sequencing data. PSI = percent spliced in. i. STRING plot demonstrating predicted protein-protein interactions of 42 shared ASEs. The PPI network was visualized via Cytoscape (v3.10.4). Yellow circle highlights the Rac1-WAVE-Arp2/3 complex node. PPI=protein-protein interaction. j. Venn diagram showing overlap between significant RBFOX2 regulated ASEs in Group 3 MB models (HDMB03/PDX7) and significantly RBFOX2-correlated exons in Group 3 MB and/or Group 4 MB patient samples, which includes core cytoskeletal genes *MYL6*, *CLSTN1* and *CTTN*. k. GO Molecular Function (GO MF) enrichment for shared significantly RBFOX2-regulated ASEs in Group 3 MB models and RBFOX2-correlated exons in Group 3 MB and Group 4 MB patient samples as shown in (j).

To confirm direct RBFOX2 dependency, we clonally expanded our CRISPR-Cas9 RBFOX2 knockout cells to remove residual RBFOX2 levels (Extended Data 6c) and subsequently re-expressed RBFOX2. Successful knockout and rescue were confirmed by immunoblotting (Fig. 5c). All seven RBFOX2-dependent splicing events showing exon exclusion in siRBFOX2 cells were fully recapitulated in the RBFOX2 knockout cells (Extended Data 6d). Importantly, RBFOX2 re-expression, in this cellular context, completely restored exon inclusion (Fig. 5d), demonstrating that these AS events are reversible and directly controlled by RBFOX2 expression. Consistent with RBFOX2-mediated growth-restrictive activity, RBFOX2 re-expression cells also reduced cell number and viability in both tumorspheres and adherent cultures (Extended Data 6e-f).

To assess whether RBFOX2-dependent AS is conserved across MB subgroups, we analyzed splicing alterations following RBFOX2 silencing in additional MB models including culture-adapted PDX7, ONS76, and culture-adapted PDX12 cells. Across all models, we identified 42 shared RBFOX2-regulated splicing events, including six of the seven candidate cytoskeletal transcript targets (*MYL6*, *ABI1*, *CLSTN1*, *MPRIP*, *CTTN* and *LIMCH1*) (Fig. 5e, Supplementary Table S10) with significant PSI reductions observed in a subgroup-independent manner (Fig.5f-h). *ENAH* also showed a significant reduction in PSI across all models but did not reach the threshold of >10% ΔPSI. Functional enrichment analysis of this AS module revealed strong association with cytoskeletal organization, cell adhesion and actin dynamics (Extended Data 7a), indicating that RBFOX2 governs a core, conserved cytoskeletal splicing program in MB development.

Protein-protein interaction (PPI) analysis of the shared RBFOX2-regulated AS module revealed a network centered on proteins associated with the RhoGTPase Rac1, a key activator of the WAVE regulatory complex (WRC)-ARP2/3 signaling pathway that drives actin polymerization and lamellipodia formation^41–45^ (Fig. 5i). The presence of *ABI1*, *MYL6*, *CTTN* and other cytoskeletal regulatory factors in this cluster suggests that RBFOX2-mediated AS converges on Rac1-dependent cytoskeletal machinery. Consistent with this model, Rac1 inhibition using the small-molecule inhibitor GYS32661 selectively suppressed expansion of RBFOX2-silenced cells with significant, but modest overall effects on cell viability (Extended Data 7b-c).

Extending these analyses to primary patient tumors, we examined bulk RNA-seq data from 480 MBs (187 Group 3 and 293 Group 4)^27^, and identified 106 shared ASEs significantly correlated with RBFOX2 expression across tumors, including *MYL6*, *CTTN* and *CLSTN1*, which were shared between all groups (Fig. 5j, Supplementary Table S11). These events were enriched for cytoskeletal/actin-associated functions and small GTPase binding (Fig. 5k). Similar enrichment was observed among OTX2-correlated AS events (Extended Data 7d-e, Supplementary Table S12). Collectively, these results demonstrate that RBFOX2 directly and reversibly governs a conserved cytoskeletal AS program across MB subgroups, with RBFOX2 loss engaging Rac1-dependent actin signaling to promote tumor cell expansion.

### RBFOX2-regulated alternative splicing events are functionally relevant

To interrogate the functional role of a RBFOX2-regulated AS event, we chose *CLSTN1*, as the specific ASE was significantly correlated with both RBFOX2 and OTX2 in MB patient tumors (Fig. 5j, Extended Data 7d). Importantly, exon 11 exhibited the most pronounced and consistent reduction in PSI across MB models following RBFOX2 silencing (Fig. 5a-b, f-h) This RBFOX2-regulated splicing event (exon 11, hg38) has previously been implicated in tumor progression^21,25^.

To directly test whether *CLSTN1* exon skipping contributes to phenotypes associated with RBFOX2 loss in our Group 3 MB models, we designed a splice-switching morpholino (Mo) to selectively induce exon 11 skipping. *CLSTN1*-Mo treatment significantly reduced PSI value relative to CTRL-Mo in HDMB03 cells (Extended Data 8a). Functionally, *CLSTN1* exon skipping induced a significant phenotypic change, shifting adhesive interactions from cell-cell contacts towards cell-substrate attachment, resulting in loss of semi-adherent cell aggregates and an increase in cell spreading and protrusive borders compared with controls (Extended Data 8b). In support of these findings, transcriptomic profiling following *CLSTN1*-Mo treatment revealed upregulation of migration-associated programs, including wound healing and chemotaxis (Extended Data 8c, Supplementary Table S13), alongside downregulation of neuronal differentiation and contractile pathways (Extended Data 8c, Supplementary Table S13). Furthermore, hallmark EMT pathways were upregulated following *CLSTN1*-Mo treatments, with concomitant reduction in hallmark myogenesis pathways (Extended Data 8d, Supplementary Table S14). These results demonstrate that inducing exon skipping of an RBFOX2-regulated splicing event phenocopies key adhesion-related and transcriptional changes associated with RBFOX2 loss, supporting its role as a downstream effector of the RBFOX2-regulated splicing program.

### RBFOX2 stratifies cell state in MB patient tumors

Our findings identify RBFOX2 as a key mediator of malignant cell-state plasticity through coordinated regulation of AS and transcriptional programs governing cycle progression, cytoskeletal dynamics and cellular plasticity. We therefore hypothesized that RBFOX2^low^ cells represent highly plastic tumor populations enriched for aggressive cellular states in MB patient tumors. To test this, we examined RBFOX2 expression across two independent published scRNA-seq datasets from MB patient tumors.

In the dataset reported by Liu *et al*., tumor cells were classified into differentiated, dividing, and quiescent cell states across both Group 3 and SHH MB^46^ (Fig. 6a-b). In Group 3 MB, stratification by RBFOX2 expression quartiles revealed significant enrichment of dividing cells within the RBFOX2^low^ (Q1) population, whereas RBFOX2^high^ (Q4) cells were enriched for quiescent and differentiated states (Fig. 6c, Extended Data 9a). Consistent with our functional RBFOX2 perturbation studies, RBFOX2^low^ Group 3 MB cells were enriched for hallmark EMT and proliferative signatures, indicative of a highly plastic transcriptional state characterized by concurrent activation of EMT-associated and tumor growth programs^9,34,35^ (Fig. 6d, Supplementary Table S15). Similar trends were observed in SHH MB tumors, with increasing RBFOX2 expression associated with a shift towards more differentiated and less proliferative states (Extended Data 9b-d, Supplementary Table S16).

**Figure 6:**
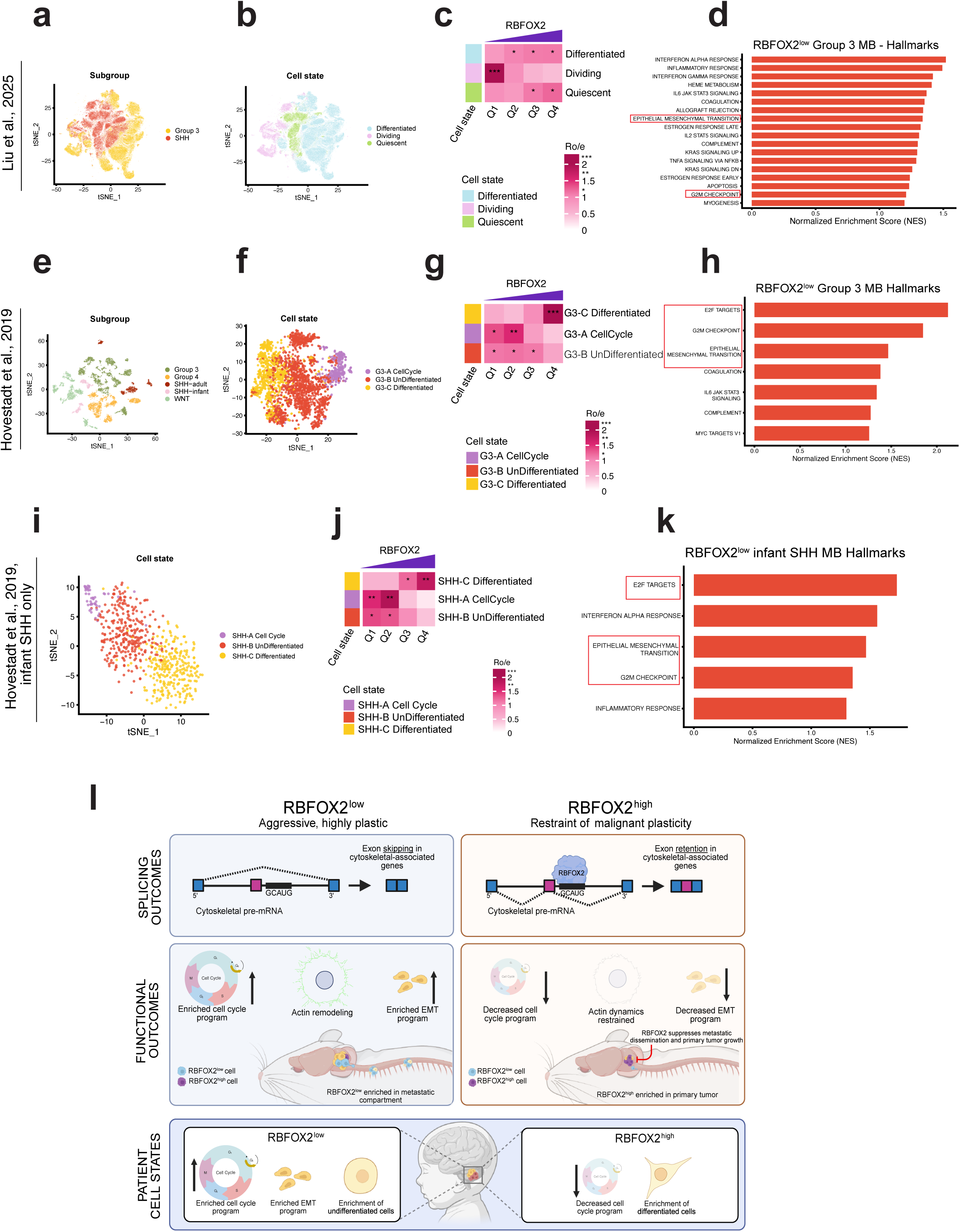
RBFOX2 defines a continuum of malignant cell plasticity in patient MB. a-b. UMAP visualization of single-cell profiles from the MB patient cohort reported by Liu et al^46^. Tumor cells are colored by MB subgroup (a) and cell states (b). c. Liu et al-derived cell state composition in Group 3 MB across RBFOX2 expression quartiles (Q1–Q4, lowest to highest) ***Ro/e >3, ** Ro/e 1.5-2. *Ro/e >1. d. Hallmark gene set enrichment analysis (GSEA) comparing RBFOX2^low^ (Q1) and RBFOX2^high^ (Q4) Group 3 MB cells in the Liu et al cohort. e-f. UMAP shows the single-cell profiles from MB patient cohorts reported by Hovestadt et al^33^. Tumor cells are colored by MB subgroups (e) and cell states (f). g. Cell state composition across RBFOX2 quartiles from lowest Q1 to highest Q4 in Group 3 MB. ***Ro/e >3, ** Ro/e 1.5-2. *Ro/e >1. G3 = Group 3 MB. h. Hallmark GSEA comparing RBFOX2^low^ (Q1) versus RBFOX2^high^ (Q4) Group 3 MB cells in the Hovestadt et al. cohort. i. UMAPs shows the single cell profiles of SHH MB infant patient dataset from the Hovestadt et al. cohort, with cells colored by cell states. j. Cell state composition across RBFOX2 expression quartiles (Q1–Q4), lowest to highest) in SHH MB infant patient dataset from the Hovestadt et al. cohort. k. Hallmark gene set enrichment analysis (GSEA) comparing RBFOX2^low^ (Q1) and RBFOX2^high^ (Q4) tumor cells in SHH MB infant patient dataset from Hovestadt et al. cohort. l. Schematic model illustrating how RBFOX2-dependent alternative splicing programs control MB cell state plasticity. Left (RBFOX2^low^): reduced RBFOX2 expression promotes widespread alternative splicing changes in cytoskeletal and cell motility regulators resulting in a highly plastic tumor cell state. This state is characterized by enhanced proliferative capacity and is enriched in metastatic cells. Right (RBFOX2^high^): elevated RBFOX2 expression redistributes RBFOX2 to downstream permissive cytoskeletal-associated exons, enforcing a splicing program that restrains malignant cell plasticity, including EMT-associated programs, undifferentiated cell states and metastatic competence. In primary patient MB (bottom), RBFOX2 expression stratifies MB tumor cells along a differentiation continuum across multiple MB subgroups, with RBFOX2^low^ states (left) linked to the most aggressive disease states while increased RBFOX2 expression (right) shows reductions in these oncogenic properties. Model created using BioRender.

We next analyzed an independent MB patient dataset including all MB subgroups reported by Hovestadt *et al*.^33^ (Fig. 6e). In Group 3 MB, the data were annotated into cell cycle, undifferentiated, and differentiated metaprograms (Fig. 6f). RBFOX2^high^ cells (Q4) showed a significant increase in the proportion of differentiated cells at the expense of undifferentiated and cycling populations (Fig. 6g, Extended Data 9e). RBFOX2^low^ Group 3 MB cells were enriched for hallmark EMT and cell cycle-related pathways (Fig. 6h, Supplementary Table S17). In SHH MB, we observed a significant increase in the proportion of differentiated cells and a reduction in the proportion of undifferentiated cells with increasing RBFOX2 expression (Extended Data 9f-h), although hallmark EMT enrichment was not observed overall in SHH RBFOX2^low^ cells (Extended Data 9i, Supplementary Table S18). However, when we restricted our analyses to infant SHH MB patient tumors, we observed patterns that are strikingly similar to Group 3 MB tumor cells with RBFOX2^low^ cells enriched for undifferentiated/cycling states as well as hallmark EMT- and cell cycle-associated programs (Fig. 6i-k, Extended Data 9, Supplementary Table S19). Collectively, these findings position RBFOX2 as a conserved determinant of malignant cell state stability across MB patient subgroups, with RBFOX2^low^ tumor cells enriched for highly plastic cellular states marked by the co-activation of undifferentiated, proliferative, and EMT-like programs.

## Discussion

Our study identifies RBFOX2 as a bifunctional regulator of malignant cell-state plasticity in MB, acting through integrated AS and transcriptional programs that converge on cytoskeletal remodeling, linage identity, and metastatic competence. Across culture-adapted models, and patient tumors, reduced RBFOX2 activity promoted highly proliferative, undifferentiated and EMT-associated cellular states, linking cytoskeletal regulation to a highly plastic, aggressive cell phenotype (Fig. 6l). However, once this high-risk state is established, additional RBFOX2 reduction does not further amplify tumor progression *in vivo*, consistent with a saturated plastic state. Notably, RBFOX2 silencing induces robust upregulation of myogenic pathways consistent with enhanced actomyosin remodeling and the non-muscle cell contractome^47^. These findings are in line with emerging evidence linking Group 3 MB myogenic transcriptional signatures with poor clinical outcomes^48^ and suggest that RBFOX2 suppresses the acquisition of contractile transcriptional programs associated with malignant Group 3 MB progression. Together, with our snRNA-seq analyses, these observations indicate that RBFOX2 constrains discrete cell state transitions.

We have uncovered a previously unrecognized antagonistic regulatory axis between OTX2 and RBFOX2 which co-regulate overlapping yet directionally opposed splicing events, particularly within genes controlling cytoskeletal dynamics. Through this antagonism, RBFOX2 limits tumor progression and is also associated with differentiated cellular identities in Group 3 MB patient tumors. While OTX2 is firmly established as a key driver of primitive oncogenic states in Group 3 MB, our findings extend this framework by positioning RBFOX2 as an antagonistic post-transcriptional regulator that tempers this oncogenic program. Mechanistically, we demonstrate that OTX2 modulates RBFOX2 activity by reshaping its RNA binding landscape. Loss of OTX2 increases RBFOX2 expression and redistributes RBFOX2 binding toward permissive intronic regions, favoring exon inclusion in accordance with established RBFOX2 positional preferences. These findings highlight OTX2 not only as a transcriptional regulator, but also as an upstream determinant of splicing factor engagement, revealing how transcriptional oncogenes can indirectly rewire AS programs that shape cellular behavior.

Mapping RBFOX2-regulated splicing activity onto the developing human cerebellum reveals a striking developmental parallel. OTX2-associated splicing patterns align with primitive RL stem cell compartments, while RBFOX2-associated splicing is repressed in these early developmental stages. The patterns suggest that OTX2-RBFOX2 antagonism reflects a conserved regulatory paradigm in human cerebellar development that is co-opted in tumorigenesis. In this framework, RBFOX2 functions as a developmental gatekeeper, restricting the persistence of undifferentiated highly plastic states.

Recent models of EMT have shifted from viewing the process as a binary switch to a continuum of reversible cellular states that endow cells with distinct phenotypic and functional properties during tumor evolution. Accumulating evidence suggests that tumor dissemination and metastatic colonization are often driven by cells occupying intermediate or hybrid EMT states that simultaneously retain proliferative capacity, stem-like features, and migratory potential^34,35,49–51^. Although MB does not arise from a classic epithelial lineage, our data parallel principles emerging from the broader EMT-plasticity field. RBFOX2 loss promotes cytoskeletal remodeling and activation of EMT-associated transcriptional programs while retaining proliferative capacity and undifferentiated cellular identities. These features are consistent with highly plastic tumor populations in which stem-like, EMT-like and proliferative programs co-exist^34,35,49–54^. In agreement with this model, RBFOX2^low^ populations identified in MB xenografts and patient samples were similarly enriched for concurrent activation of undifferentiated, cycling, and EMT-associated programs. Furthermore, RBFOX2 overexpression reduced leptomeningeal dissemination *in vivo*, linking suppression of these plastic transcriptional states with reduced metastatic competence. These findings position RBFOX2 as a determinant of cellular state stability, limiting the emergence of highly plastic Group 3 MB cell states. Importantly, both transcriptional and post-transcriptional gene regulation, specifically AS, have been shown to regulate cell state along the EMT continuum^34,51^. Our findings suggest that RBFOX2 coordinates transcriptional and AS programs that converge on cytoskeletal dynamics to regulate malignant cellular plasticity in Group 3 MB.

AS has emerged as an important post-transcriptional regulator of cell identity, stemness and state transitions across the EMT continuum^40,51^. We identified a conserved RBFOX2-regulated cytoskeletal splicing module shared across Group 3 and/or SHH MB models. This module includes key regulators of actin remodeling and cell motility such as *MYL6*, *ABI1*, *CLSTN1*, *MPRIP*, *ENAH*, *CTTN* and *LIMCH1*. Functional rescue experiments demonstrate that these splicing events are directly dependent on RBFOX2, establishing a causal link between RBFOX2 activity and cytoskeletal isoform selection. Network analysis further revealed that this splicing module converges on a Rac-dependent WAVE-Arp2/3 signaling axis, a central diver of actin dynamics and cell migration. Notably, profiling of primary Group 3 MB patient tumors identified Rho-GTPase signaling as a key vulnerability in aggressive disease^55^.

Together, our findings establish RBFOX2 as a key cell-state stabilizer that constrains malignant cell plasticity through the coordinated transcriptional and post-transcriptional control of cytoskeletal dynamics. RBFOX2 limits the emergence of highly proliferative, undifferentiated and EMT-like tumor cell states that are linked to metastatic competence. More broadly, our work highlights how antagonistic interactions between oncogenic transcription factors and splicing regulators sculpt developmental identity and cellular plasticity in MB, revealing AS as a critical regulatory layer with potential therapeutic implications.

## Methods

### Cell lines and patient-derived xenograft (PDX) *in vitro* culturing conditions

Group 3 MB cell lines HDMB03^56^ and MB3W1^37^ were generously provided by the Milde and Wölfl laboratories, respectively. D283 cells lines were purchased from American Type Culture Collection (ATCC, Rockville, MD, USA), HEK293T was purchased from ATCC and ONS76 was provided by Dr. Mark Remke. HDMB03 and MB3W1 tumorspheres were cultured in StemPro NSC and stem cell media^5,8^ (DMEM/F12, 2% B27, 1% MEM vitamin solution, 20 ng/mL bFGF, 20 ng/mL EGF, 40 U/ml penicillin, and 40 μg/ml streptomycin), respectively. D283 tumorspheres were cultured in DMEM/F12, 2% B27, 1% N2, 40ng/mL bFGF, 40ng/mL EGF^5^. HEK293T and ONS76 were cultivated in Dulbecco’s Modified Eagle’s medium (DMEM) + GlutaMAX-I high glucose medium (Thermo Fisher Scientific) supplemented with 10% fetal bovine serum (FBS) and 1% penicillin/streptomycin. RPMI media supplemented with 10% FBS was used for HDMB03 adherent culture experiments. Patient-derived xenograft (PDXs) ICN-MB-PDX12 (PDX12), ICN-MB-PDX7 (PDX7) were generated from primary human MBs obtained from surgical resection performed at the Children’s Necker Hospital in Paris (France). Human samples were obtained with informed consent of patients, and all experimental procedures were performed following guidelines from the Institutional Review Board at Necker Hospital, Paris, France. Once established, PDX models were maintained *in vivo* by serial propagations. For *in vitro* cultures, PDX tumors were dissociated in Neurobasal Media containing 1 mg/ml DNaseI (Worthington Biochemicals), 2.5 mg/ml Collagenase P and 2.5 mg/ml Collagenase/dispase (Roche), B27 supplement minus vitamin A, and N2 supplement, and then they were separated by centrifugation on a Percoll density gradient (Merck). Purified PDX12 cells were cultured in Neurobasal Media supplemented with B27 supplement minus vitamin A, 0.01% BSA solution, 1% 1000X N-Acetyl Cysteine (Sigma-Aldrich) and 1% d+-Glucose solution 45% (Sigma-Aldrich), 1% Pen–Strep and 1% l-Glutamine. Purified PDX7s were grown in Neurobasal Media supplemented with B27 supplement minus vitamin A, N2 supplement, BSA solution, 1% Pen–Strep and 1% l-Glutamine, 20ng/ml human bFGF, and 20ng/ml human EGF. Cells were incubated at 37°C, 5% CO_2_ for 3 or 5 days depending on downstream assays. Cells prepared for live cell counts and viability assays were plated at 1.5X10^5^ cells/well in 6 well ultra-low attachment plates (3D culture, Corning) or standard tissue culture plates (adherent).

### siRNA silencing

Silencing of OTX2 and LASR gene (RBFOX2, HNRNPC, and HNRNPM) expression was performed in Group 3 MB cells using silencer select siRNAs while a scramble sequence served as a negative control (Invitrogen). Knockdowns were performed at 30 nM concentration. Silencing was performed using Lipofectamine RNAiMAX Transfection Reagent (Invitrogen). Sequences for siRNA are provided in Supplementary Table S20. For combination experiments with siRNAs and GYS32661, HDMB03 cells were seeded at a density of 1.5 × 10^5^ cells per well in 6-well plates with three technical replicates per condition. Cells were transfected the following day with either RBFOX2-targeting siRNA (Silencer Select siRNA, s24049) or non-targeting control siRNA (Silencer Select Negative Control No. 1 siRNA, 4390843) at a final concentration of 30 nM using Lipofectamine RNAiMAX in Opti-MEM Reduced Serum Medium according to the manufacturer’s instructions. Eight hours post-transfection, cells were treated with either 20 μM GYS32661 or 0.2% DMSO vehicle control and maintained for 72 hours. Following treatment, cells were dissociated, harvested, and quantified for total cell number and viability.

### Tumorsphere size assay

For RBFOX2 knockdowns, HDMB03 cells were seeded at a density of 2500 cells per well in 24-well ultralow attachment plates (Corning). RBFOX2 silencing was performed as described above. Three wells were seeded per condition in each independent experiment. For the RBFOX2 overexpression, HDMB03-Luc-RBFOX2 OE cells were seeded at a density of 100 cells per well into ultra-low attachment 96-well round-bottom plates (Corning Costar). Twenty wells (technical replicates) were seeded per experimental group. Tumorspheres were imaged 72 h post-seeding using an EVOS XL Core microscope, and sphere area was quantified using FIJI (ImageJ). Individual sphere measurements were treated as technical replicates nested within three independent biological experiments. Statistical comparisons between conditions were performed using a nested unpaired t-test in GraphPad Prism, with the biological experiment defined as the nesting factor.

### Lentiviral production and infection

Lentiviral particles were generated in HEK293T cells by transiently transfecting the packaging plasmid psPAX2 and the envelope plasmid pMD2.G with lentiviral vectors (shSCR #VB010000-9526zpu), shRBFOX2-1 #VB900089-8432ckt, shRBFOX2-2 #VB900089-8438vxu, purchased from Vector Builder) using X-tremeGENE™ 9 DNA transfection Reagent. Supernatants were harvested at 48 h and 72 h and used to infect cells for 72 h. After infection, transduced cells were selected using 1-2 μg/mL Puromycin dihydrochloride (TOKU-E EU N.V).

For these infections, cells were seeded at 40% confluency and cultured in a 1:1 mixture of fresh media and viral suspension. For RBFOX2 knockdown, we used the combination of shRNA lentiviral vectors: shRBFOX2-1 and shRBFOX2-2.

### Morpholino treatment

HDMB03 cell cultures were treated with a *CLSTN1* splice-blocking morpholino (Gene Tools) at final concentrations of 1 µM and 2 µM for 48 hours. Morpholino sequence information is provided in Supplementary Table S20. Before harvest, cells were imaged to assess morphology and adhesion characteristics. Imaging was performed at 20x magnification using an EVOS XL Core microscope. Following the 48-hour treatment period, cultures were dissociated using StemPro Accutase. Total RNA was isolated using the Norgen Biotek Total RNA Purification Kit following the manufacturer’s protocol. For RNA sequencing, 2 μM CLSTN1-Mo samples were used. Primers for exon skipping validation are provided in Supplementary Table S20.

### CRISPR-Cas9-based RBFOX2 knockout

For CRISPR–Cas9-directed knockout, two single guide RNAs (sgRNAs) targeting exons 2 and 3 of the RBFOX2 gene, along with a non-targeting control (NTC) sgRNA, were designed, synthesized, and cloned into the LentiCRISPR v2 vector (Addgene #52961) as previously described^21^. Successful integration of the guide sequences was confirmed by Sanger sequencing with the U6 forward primer. Lentiviral particles were prepared by the ATC Gene Vector Core at Baylor College of Medicine at a titer of more than 1 × 10^8 TU/mL. Briefly, lentiviral particles were produced by co-transfecting HEK293T cells with a second-generation packaging plasmid and either sgRBFOX2 LentiCRISPRV2 or sgCTRL LentiCRISPRV2 using iMFectin transfection reagent (GenDepot). HDMB03 cells were transduced with viruses at an MOI of 3. Cells were selected with puromycin (2 μg ml-1, ThermoFisher) 48 hours post-transduction for 7 days, and followed by further expansion using single-cell sorting on a Sony MA900 (Cytometry and Cell Sorting Core, Baylor College of Medicine, TX, USA) into 96-well plates to isolate pure RBFOX2 KO colonies. Individual clones were expanded. Immunoblot analysis and Sanger sequencing were performed to confirm loss of RBFOX2 expression. CRISPR sgRNA sequences and primers for knockout validation are provided in Supplementary Table S20.

### RBFOX2 overexpression

To overexpress RBFOX2, lentiviral particles containing RBFOX2 cDNA, NM_001031695 (Cat#RC206846) or negative control shuttled in pLenti-C-MYC-DDK-P2A-BSD (Cat#PS100103) vector were purchased from Origene. HDMB03 LV-CMV5-Luc2-GFP and HDMB03 sgRBFOX2 KO cells were transduced at an MOI of 1 and 5. Cells were selected with Blasticidin (25 μg/mL) after 48 hours of post-transduction. Immunoblot analysis was performed to confirm overexpression.

### Immunoblotting

Proteins ranging from 10-50 μg were measured using the BCA Protein Assay kit (Pierce) and resolved on Bolt™ Bis-Tris Plus Gels (Thermo Scientific). Proteins were transferred onto nitrocellulose Power Blotter Transfer Stacks using the Power Blotter Transfer system (Invitrogen). Primary antibodies utilized are summarized in Supplementary Table S20. Images were acquired using the Fusion FX Vilber Lourmat chemiluminescence imaging system (Vilber).

### Immunofluorescence and phalloidin staining

Cells were grown on glass coverslips in 24-well plates. To preserve the dynamic actin structure, cells were removed from the incubator and immediately subjected to mild permeabilization and fixation with 0.1% Triton X-100 and 4% PFA for 5 minutes. The solution was replaced with 4% PFA alone, and fixed for 10 min. Following PBS washing 3 times, coverslips were blocked in 1% bovine serum albumin (BSA) in PBS for 1 hr. Filamentous actin (F-actin) was stained with phalloidin labelling probes (488-Fisher) at a 1:200 dilution for 30 minutes at room temperature in the dark. After phalloidin staining, coverslips were washed 3 times with PBS-T (0.1 % Tween 20). Coverslips were counterstained with Hoechst 33342 at 1:1000 for 10 minutes, washed 3 times with PBS-T, and mounted using a Prolong Diamond Antifade Mountant (Invitrogen). Slides were stored in the dark at 4°C until imaging. Images were acquired using a Nikon A1R-s Confocal Microscope (Integrated Microscopy Core, Baylor College of Medicine) at identical settings across samples. Images were further processed and analyzed using Fiji (ImageJ). For each condition, cells were manually traced using the Freehand Selection tool. For each ROI, the following parameters were extracted using Analyze → Measure: Integrated density (IntDen) intensity (phalloidin channel), area, and circularity. Data were analyzed in GraphPad Prism. Groups were compared using a Mann-Whitney test.

### RT-PCR

Total RNA was isolated using an RNA Purification Kit (Norgen Biotek). For cDNA synthesis, 2 μg of total RNA was reverse-transcribed with the SuperScript III First-Strand Synthesis kit (Life Technologies). PCR amplification was performed with MiniAmp Plus Thermal Cycler (Applied Biosystems, Thermo Fisher Scientific) as previously described^8^. PCR products were run on a 2% ultrapure agarose gel (Invitrogen, Life Technologies), stained with EZ-vision Bluelight DNA dye, and visualized using a Fusion FX Vilber Lourmat chemiluminescence imaging system. At least three splicing PCRs were performed for each cell line and target exon. Primer sequences are listed in Supplementary Table S20.

### RNA extraction, bulk RNA-sequencing, and analysis of alternative splicing events in silenced RBFOX2 cells

For siRBFOX2, siHNRNPM and siHNRNPC-treated HDMB03 cells, RNA was extracted using an RNA purification kit (Norgen Biotek) per manufacturer’s instructions. RNA isolation for RNA sequencing of shRBFOX2-depleted PDX7, ONS76, and PDX12 cell pellets was performed using RNeasy Plus Mini Kit (Qiagen) as per the manufacturer’s protocol. RNA quality was assessed with the Agilent RNA 6000 Nano kit (Agilent) on a Bioanalyzer (Agilent). Total RNA (1 μg) was processed into barcoded libraries using the TruSeq Stranded RNA LT Kit (low-throughput protocol, Illumina). Libraries were sequenced on the Illumina NovaSeq platform using a paired-end read length of 100 base pairs (NovaSeqX - 1.5B-200 - 1U - PE100). Subsequent demultiplexing of the raw BCL files yielded FASTQ files containing the sequencing data.

RNA-seq reads were mapped to the human genome using the hg38 reference with the STAR aligner (version 2.7.11b) in two-pass mode, with GENCODE v45 annotations. SAMtools (v1.9) and Picard MarkDuplicates (v2.18.14) were used to coordinate-sort BAM files and flag duplicate reads, respectively, and low-mapping-quality reads (MAPQ < 20) were subsequently removed. rMATS (version 4.3.0)^57^, an event-based tool, was used to identify differentially spliced events between control and knockdown conditions for OTX2, RBFOX2, HNRNPM and HNRNPC, using GENCODE v45 annotations. Five distinct alternative splicing events were quantified: skipped exons (SE), mutually exclusive exons (MXE), alternative 5’ splice sites (A5SS), alternative 3’ splice sites (A3SS), and retained introns (RI). Briefly, rMATS features a count-based model that calculates PSI values across samples using spliced reads and reads that map to the exon body. To call statistically significant splicing events, each splicing event had to: (1) have a |ΔPSI| of >10%; (2) have a minimum support of 15 unique reads; and (3) have a FDR below 0.05.

Alignment parameters: --outFilterMultimapNmax 10 --alignSJoverhangMin 5 --alignMatesGapMax 1000000 --alignIntronMax 1000000 --chimSegmentReadGapMax 0 --alignSJDBoverhangMin 3 --alignSJstitchMismatchNmax 5 -1 5 5 --outSAMmultNmax 20 --chimSegmentMin 12 --chimJunctionOverhangMin 12 --outBAMsortingThreadN 8 --twopassMode Basic rMATS parameters: --readLength 100 --variable-read-length --libType fr-firststrand -t paired --cstat 0.0001 --allow-clipping.

### Correlations between OTX2- and RBFOX2-regulated exons with Group3/Group4 patient samples

To explore OTX2- and RBFOX2-regulated exons in MB, we re-analyzed previously published bulk RNA-seq results for Group 3 and Group 4 MB subgroups. Briefly, we kept exons that were expressed in at least 20% of samples and having a max(PSI) – min(PSI) ≥ 0.2 and excluded samples having more than µ + 3σ of non-detected exons. We performed a correlation-based analysis (cor.test function in R) between OTX2 and RBFOX2 VST-normalized expression and PSI values. Correlation coefficients and associated p-values were extracted for all complete exon–gene pairs, and p-values were adjusted for multiple testing using the Benjamini–Hochberg FDR method. Splicing events with an adjusted p-value of < 0.01 were retained as significantly correlated with gene expression.

### Motif Enrichment Analysis

Motif enrichment analysis was performed using rMAPS2^58^ to identify RNA-binding proteins motifs enriched around differentially spliced events. Briefly, regulated splicing events identified from rMATS output were separated into upregulated and downregulated sets according to the direction of ΔPSI change and compared against a background set of SE events annotated in GENCODE v45 that were detected in the dataset but did not belong to the regulated splicing signature. When the background exceeded 15,000 events, a random subsample of 15,000 events was used. rMAPS2 examines adjacent intronic flanking sequences in upstream and downstream regions (±250 nt) relative to the regulated exon and computes enrichment scores independently for each region. Enrichment p-values for all RBPs were subsequently extracted from rMAPS2 output tables, transformed as −log10(p-value), and ranked by statistical significance, and the top 15 enriched RBPs were retained for visualization.

### Splicing signatures of OTX2 and RBFOX2 mapped onto cerebellar data

To investigate OTX2- and RBFOX2-regulated exons in the human cerebellar developmental context, we used previously published cerebellar bulk RNA-seq data for the rhombic-lip (RL^VZ^ (n=7) and RL^SVZ^ (n=8) at post-conception weeks (PCWs) 9 to 22) and EGL (at PCW 14 to 22; n=15). For each splicing event shared between the developmental cerebellar dataset and the OTX2- and RBFOX2-regulated exon signatures, PSI values were sign-adjusted according to the direction of regulation (ΔPSI sign) and centered across samples to account for baseline differences in global splicing levels. For each sample, the splicing signature score was defined as the mean of all centered, sign-adjusted PSI values across shared signature events. Owing to centering, scores reflect relative deviations in splicing activity; a score of 0.1 corresponds to an average shift of 10% in exon inclusion or exclusion in the direction of the signature relatively to the cohort mean.

To assess the statistical significance of the splicing signature scores, we employed a permutation-based approach to determine whether the observed scores differed from what would be expected by chance. For each condition, a null distribution was generated by computing scores from 1000 randomly drawn sets of splicing events matching the mean size of the splicing signatures. The observed scores were then compared against the null distributions to derive empirical p-values adjusted for multiple testing using the Benjamini–Hochberg FDR method. Significance levels were denoted as follows: *** FDR < 0.001, ** FDR < 0.01, and * FDR < 0.05.

### Enhanced Crosslinking and Immunoprecipitation (eCLIP)

eCLIP was performed as previously described^59^. Briefly, two replicates each of siCTRL or siOTX2 HDMB03 tumorspheres were UV crosslinked (400 mJ/cm2 at 254 nM), pelleted, and snap frozen in liquid nitrogen. Pelleted cells were then lysed in eCLIP lysis buffer, treated with RNase I to fragment RNA, and subjected to immunoprecipitation with an antibody against RBFOX2, HNRNPM or OTX2 and sheep anti-rabbit secondary Dynabeads (ThermoFisher) (Supplementary Table S20). After washes, dephosphorylation, and RNA adapter ligation, immunoprecipitation and paired input samples were run on 4-12% Bis-Tris SDS-PAGE gels and transferred to a nitrocellulose membrane. The region from 50 to 125 kDa was isolated for RBFOX2 and HNRNPM eCLIP, and the region from 32-107 kDa was isolated for OTX2 eCLIP. After Proteinase K treatment, RNA was isolated, reverse transcribed (Superscript IV, ThermoFisher), a 3’ DNA adapter was ligated (T4 RNA Ligase, NEB), and samples were PCR amplified and sequenced on the Illumina NovaSeq XP platform.

Processing and analysis of eCLIP data were performed as previously described^59^. Briefly, reads were adapter trimmed (cutadapt v3.4), mapped (STAR v 2.4.0j) against repeat elements (with mapped reads removed from further analysis), and then mapped (STAR v2.4.0j) against the human genome (hg38) with only uniquely mapping reads retained. PCR duplicates were then removed (Umi-tools v1.1.1), and peak calling (CLIPPER v2.0.0) and input normalization were performed as previously described^59^. Splicing maps were generated combining differential exon lists from OTX2, RBFOX2, or HNRNPM knockdown RNA-seq and RBFOX2 or HNRNPM eCLIP data using the previously described RBP-Maps software [[PMID 30413564]].

### scRNA-sequencing on Group 3 MB tumorspheres

HDMB03 tumorspheres were transfected with siRNAs (Silencer Select negative control or RBFOX2 siRNA, s24049, Thermo Fisher Scientific (Supplementary Table S20), or transduced with control particles or RBFOX2 overexpression constructs prior to single-cell RNA sequencing. Tumorspheres were dissociated into single-cell suspensions by gentle trituration, followed by centrifugation (250 × g, 5 min, 4 °C), resuspension in ice-cold PBS supplemented with 10% FBS, and filtration through a 40 μm cell strainer. Cells were pelleted (150 × g, 3 min, 4 °C), washed twice in ice-cold PBS containing 0.04% BSA, and filtered prior to sorting. Viable single cells were enriched by fluorescence-activated cell sorting using DAPI exclusion to minimize debris and processed immediately. Single-cell libraries were generated using the 10x Genomics Chromium Single Cell 3’ v4 Gene Expression kit, targeting ∼20,000 cells per sample. Libraries were pooled and sequenced on an Illumina NovaSeq X platform (10B flow cell; paired-end 151 bp), yielding ∼200 million reads per sample.

### Data integration, normalization, batch correction, and clustering

Raw sequencing data were demultiplexed, aligned to the GRCh38 human reference genome, and processed to generate barcode–gene UMI matrices using Cell Ranger (v9.0.1, 10x Genomics). The resulting UMI matrices were subsequently analyzed in R (v4.4.1) using Seurat (v5.3.1)^60^. Additional quality control was performed to remove low-quality cells. Cells were excluded if they met any of the following criteria: (1) fewer than 200 detected genes or a number of detected genes exceeding five standard deviations above the sample median; (2) total UMI counts exceeding five standard deviations above the sample median; or (3) a mitochondrial transcript fraction greater than 10%.

Datasets were processed using Seurat (v5.3.1)^60^ in R (v4.4.1). In total, 72,173 cells passed QC and were retained for downstream analyses. For initial preprocessing, gene expression matrices were normalized to a uniform library size of 10,000 counts per nucleus and log-transformed using Seurat’s NormalizeData function. A total of 3,000 highly variable genes (HVGs) were identified using FindVariableFeatures. Data were subsequently scaled with ScaleData, followed by principal component analysis (PCA) using RunPCA. Graph-based clustering was performed using FindNeighbors (pc.num = 1:30) and FindClusters (resolution = 0.1), and dimensionality reduction for visualization was conducted using Uniform Manifold Approximation and Projection (UMAP)^61^ via RunUMAP with default parameter.

To remove batch effects while preserving biological signal, we applied SCTransform-based normalization SCTransform. Batch correction was then performed on the SCTransform (SCT) assay using R package Harmony (v1.2.4)^62^, with sample identity specified as the integration variable. Downstream dimensionality reduction, neighborhood graph construction, clustering, and UMAP embedding were computed using Harmony-corrected embeddings.

Three meta-programs for Group 3 MB (G3 MB), curated from Hovestadt et al.^33^, were used to annotate cells as G3-A (cell cycle), G3-B (undifferentiated), or G3-C (differentiated) tumor states. Meta-program scores were computed using Seurat’s *AddModuleScore* function (v5.3.1), and each cell was assigned to the meta-program with the highest score. Cells were further annotated for proliferative status (MKI67⁺/⁻) based on MKI67 expression. Cell-cycle scores were calculated using Seurat’s *CellCycleScoring* function (v5.3.1), and cells were assigned to cell-cycle phases according to these scores.

To quantify group-level prevalence of cell types, we calculated an observed-to-expected ratio (Ro/e), defined as

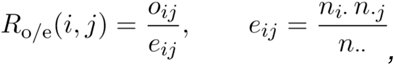

where *o_ij_* is the observed number of cells of cell type *i* in group *j* and *e_ij_* is the corresponding expected number of cells under independence. Here, *n_i_*_·_ = ^P^*_j_ o_ij_* denotes the total number of cells of cell type *i* across all groups (row total), *n*_·*j*_ = ^P^*_i_ o_ij_* denotes the total number of cells in group *j* across all cell types (column total), and *n*_··_ = ^P^*_i,j_ o_ij_* denotes the total number of cells in the dataset (grand total). Under the null hypothesis that cell type and group are independent, the expected number of cells of type *i* in group *j* is computed as *e_ij_* = (*n_i_*_·_*n*_·*j*_)*/n*_··_. A Ro/e value greater than one indicates the enrichment of a cell type in a specific group. Enrichment of cell type *i* in group *j* was defined as *R*_o*/*e_(*i,j*) *>* 1, consistent with established Ro/e analyses^63–66^. For visualization, Ro/e matrices (cell types by groups) were displayed as heatmaps using the ComplexHeatmap R package^67,68^

### Differential gene expression and pathway enrichment

Differentially expressed genes (DEGs) for each comparison were identified using Seurat’s FindMarkers function (only.pos = F) and ranked by log fold change. Gene set enrichment analysis (GSEA)^69^ was performed using R package fgsea (v1.32.4)^70^. Gene sets were obtained from the Molecular Signatures Database (MSigDB) via the msigdbr package (v25.1.1)^71^, including Hallmark gene sets (H category) and Gene Ontology Biological Process (C5: BP), restricted to Homo Sapiens. Pathways were considered significant if they had an absolute normalized enrichment score (|NES|) > 1 and an adjusted p-value < 0.05.

### Group 3 and SHH MB patient sample data analyses

We collected two published datasets which include Group 3 and SHH MB samples: (1) scRNA-seq and snRNA-seq datasets reported by Liu *et al*^46^. (2) Smart-seq2 dataset reported by Hovestadt *et al*^33^.

The Liu *et al*^46^ data were stratified by molecular subgroup for cancer subgroup-specific analyses. Each cancer subgroup was processed independently following the same analytical pipeline described below. Three gene signatures curated from Liu *et al*^46^ were used to annotate tumor cells as differentiated, dividing or quiescent. Signature scores were computed using Seurat’s AddModuleScore (v5.3.1), and each cell was assigned to the signature with the highest score.

Datasets were divided by cancer subtypes, and cells within each dataset were ranked by normalized RBFOX2 expression and stratified into quartiles: Q1 (lowest), Q2, Q3 and Q4 (highest). To evaluate compositional shifts across RBFOX2 expression levels, we calculated the relative proportions of annotated cell types (by signature assignment) within each quartile. Trends from Q1 to Q4 were assessed to determine whether increasing RBFOX2 expression was associated with cellular distribution. For downstream transcriptional comparisons, Q1 cells were defined as the RBFOX2^low^ group and Q4 cells as the RBFOX2^high^ group to maximize contrast between expression extremes. Differential gene expression and pathway enrichment analysis between RBFOX2^low^ and RBFOX2^high^ cells was performed. Ro/e analysis was used to quantify group-level enrichment of cell types as defined above, with enrichment of a cell type in a group defined as *R*_o*/*e_ *>* 1.

Differentially expressed genes (DEGs) between Q1 (RBFOX2^low^) and Q4 (RBFOX2^high^) were identified using Seurat’s FindMarkers function. Pathway-level enrichment was assessed using the fgsea package (v1.32.4)^70^ on pre-ranked gene lists derived from differential expression results. The same MSigDB gene set collections (Hallmark, GO BP, and GO MF) were applied.

The Hovestadt *et al*^33^ cohort comprised Smart-seq2 dataset representing all four MB subgroups: WNT, SHH, Group 3 and Group 4 MB. Only SHH and Group 3 MB subgroups were included in subsequent analyses. Data were stratified by molecular subgroup, and each subgroup was processed independently as described below. Three Group 3 MB and SHH MB meta-programs were curated from Hovestadt *et al*^33^ and used to annotate cells as G3-A cell cycle, G3-B undifferentiated, G3-C differentiated, SHH-A cell cycle, SHH-B undifferentiated or SHH-C differentiated tumor states. Meta-program scores were computed using Seurat’s AddModuleScore (v5.3.1), and each cell was assigned to the highest-scoring meta-program. To validate RBFOX2 perturbation-associated transcriptional programs, we applied the same in silico stratification and analyses strategies as described above for the Liu et a., dataset.

### Mouse housing and husbandry

All animal procedures were approved by the Baylor College of Medicine Institutional Animal Care and Use Committee (AN9172) and conducted in the Feigin Center specific-pathogen-free vivarium. NSG mice (7-9 weeks at implantation) were housed under sterile conditions with autoclaved cage setups and sterile water and provided with standard environmental enrichment (Enviropacks). Following stereotactic brain surgery, animals were singly housed until the incision and injection sites healed. Seven days later, after autoclips were removed, they were returned to group housing. During the immediate post-operative period, NapaNectar gel packs were placed on the cage floor as needed to support hydration and recovery, in addition to standard chow and water provided ad libitum.

### Intracerebellar implantation and tissue collection

Seven- to nine-week-old NSG mice were anesthetized with isoflurane and placed in a stereotaxic apparatus to stabilize the skull. A midline scalp incision was made, a burr hole was drilled over the cerebellum, and injections were performed using stereotaxic coordinates referenced to lambda (1.5 mm anterior-posterior, 2.5 mm dorsal-ventral). For all experiments, HDMB03 cells and their derivatives were injected at doses of 1.5×10³ or 1.5×10⁴ cells in 5 µl sterile PBS using a 10 μl Hamilton syringe fitted with a 30-gauge needle; MB3W1 cells were implanted using the same stereotactic approach at a dose of 5.0×10⁴ cells in 5 µl PBS. Tumor-bearing mice were monitored by bioluminescence imaging on days 15 and 19 post-implantation to assess primary tumor growth and leptomeningeal dissemination. At humane endpoint or defined experimental time points, animals were deeply anesthetized and transcardially perfused with saline, followed by 10% formalin. Brains and spinal cords were harvested, with a subset of tissues paraffin-embedded for histology and immunohistochemistry. Cerebellar tumors and spinal leptomeninges from additional animals were snap frozen and processed for nuclei isolation followed by single-nuclei RNA sequencing.

### Bioluminescence imaging (BLI) and analysis

Mice were anesthetized with 4% isoflurane in an induction chamber and then injected intraperitoneally with D-Luciferin at 150 mg kg⁻¹ (BD, BDB556888). Five minutes after D-Luciferin administration, animals were imaged using the IVIS Lumina III system (PerkinElmer), and images were acquired under identical settings for all groups. Bioluminescent signals were quantified using Living Image Software v4.7.3 (PerkinElmer) to obtain total flux (photons s⁻¹) from defined regions of interest.

### Immunohistochemistry

Following deparaffinization and rehydration of FFPE xenograft sections, heat-induced antigen retrieval was performed in citrate buffer (pH 6.0) using a BioCare Medical pressure-cooker system, which reached 121 °C at high pressure for 6 min and was then allowed to cool to 80 °C approximately 45 min. Slides were washed and incubated for 10 min in an endogenous peroxidase block, followed by additional washes in 1×PBS. Sections were blocked with 3% normal lamb serum and incubated overnight at 4 °C with a human-specific anti-mitochondria antibody (Supplementary Table S20). Slides were washed and incubated with a biotinylated secondary antibody (1:1000) for 2 h at room temperature, followed by Streptavidin–HRP (Vectastain ABC kit, Vector Laboratories, PK-6100) for 30 min in 1× PBS. Signal was developed using ImmPACT DAB (Vector Laboratories, SK-4105), and nuclei were counterstained with hematoxylin, after which sections were dehydrated and coverslipped with VectaMount mounting medium (Vector Laboratories, ZL-0823).

### Nuclei isolation and 10x Genomics library preparation for snRNA-seq

Cerebellar primary tumors and leptomeningeal layers from MB3W1-engrafted NSG mice were microdissected on ice, snap-frozen, and stored at −80 °C until processing. Nuclei were isolated from frozen primary tumor and leptomeningeal tissues using the Chromium Nuclei Isolation Kit (10x Genomics; User Guide CG000505, Rev. A) according to the manufacturer’s instructions. Briefly, 30–50 mg of frozen tissue was homogenized in Lysis Buffer with a plastic pestle on ice, passed through a Nuclei Isolation Column, and the flow-through containing nuclei was subjected to debris-removal and wash steps before final resuspension in Resuspension Buffer and counting by Trypan blue exclusion. After nuclei isolation, nuclei from leptomeningeal metastases from four mice were pooled into a single leptomeningeal sample. These pooled nuclei suspensions along with nuclei from a corresponding primary cerebellar tumor sample were submitted to Admera Health (South Plainfield, NJ), where 3′ single-nucleus gene-expression libraries were prepared using the Chromium GEM-X Single Cell 3′ v4 reagents (Chip, Gel Bead, GEM, and Library Construction Kit C; 10x Genomics). Samples were then sequenced on an Illumina platform using standard short-read paired-end settings.

### snRNA seq data on leptomeningeal metastasis and primary tumor cells

Raw sequencing data were demultiplexed, aligned to the human (GRCh38) and mouse (GRCm39) reference genomes, and processed to generate barcode–gene UMI count matrices using Cell Ranger (v9.0.1; 10x Genomics). The resulting UMI matrices were analyzed in R (v4.4.1) using Seurat (v5.3.1)^60^. For downstream analyses, barcodes were retained as human-derived cells only if they met both of the following criteria: (1) the barcode was classified as human (GRCh38) by Cell Ranger and (2) more than 80% of total UMI counts were derived from the human genome.

Additional quality control was performed to remove low-quality nuclei. Nuclei were excluded if they met any of the following criteria: (1) fewer than 200 detected genes or a number of detected genes exceeding five standard deviations above the sample median; (2) total UMI counts exceeding five standard deviations above the sample median; or (3) a mitochondrial transcript fraction greater than 10%. After quality control filtering, datasets were processed using Seurat (v5.3.1)^60^ in R (v4.4.1). In total, 9,238 nuclei passed QC and were retained for downstream analyses.

For initial preprocessing, gene expression matrices were normalized to a uniform library size of 10,000 counts per nucleus and log-transformed using Seurat’s NormalizeData function. A total of 3,000 highly variable genes (HVGs) were identified using FindVariableFeatures. Data were subsequently scaled with ScaleData, followed by principal component analysis (PCA) using RunPCA. Graphbased clustering was performed using FindNeighbors (pc.num = 1:30) and FindClusters, and dimensionality reduction for visualization was conducted using Uniform Manifold Approximation and Projection (UMAP)^61^ via RunUMAP with default parameter.

To remove batch effect while preserving biological signal, we applied SCTransform-based normalization SCTransform. Batch correction was then performed on the SCTransform (SCT) assay using R package Harmony (v1.2.4)^62^, with sample identity specified as the integration variable. Downstream dimensionality reduction, neighborhood graph construction, clustering, and UMAP embedding were computed using Harmony-corrected embeddings.

The three meta-programs for Group 3 MB were curated from Hovestadt *et al*^33^ and used to annotate cells as G3-A cell cycle, G3-B undifferentiated or G3-C differentiated tumor states. Meta-program scores were computed using Seurat’s AddModuleScore (v5.3.1), and each cell was assigned to the highest-scoring meta-program. To validate RBFOX2 perturbation-associated transcriptional programs, we applied the same in silico stratification and analyses strategies as described above for the Liu et a., dataset. The same analyses were used to compare primary MB and spinal cord metastases. Ro/e analysis was also used to quantify group-level enrichment of cell types, with enrichment of a cell type in a group defined as *R*_o*/*e_ *>* 1. Differentially expressed genes (DEGs) and pathway-level enrichment were assessed as described above for patient sample datasets. The same analyses were used to compare primary MB and spinal cord metastases.

### RNA-sequencing on morpholino-treated samples

HDMB03 cells were seeded at 1.5X10^5^ cells per well in adherent culture (five wells per condition) and treated with 2 μM Vivo-standard control morpholino or a CLSTN1 morpholino. Following 48 h treatment, cells were harvested, dissociated, and pelleted for RNA extraction. Total RNA was isolated using the Norgen Biotek Total RNA Purification Kit with on-column DNase I treatment. RNA quality was assessed prior to library preparation, and libraries were generated using NEBNext rRNA depletion and Ultra II directional RNA library preparation kits. Libraries were sequenced on an Illumina NovaSeq X platform (25B flow cell; paired-end 151 bp), targeting ∼100 million reads per.

Raw sequencing reads were first subjected to quality assessment using fastqc (v0.12.1). Adapter sequences and low-quality bases were trimmed using fastp (v1.0.1) with default parameters. Post trimming read quality was re-evaluated using fastqc to confirm removal of adapters and improvement of per-base sequence quality. Trimmed reads were aligned to the human reference genome (GRCh38.p14; GENCODE release v49^72^) using STAR (v2.7.11b) in two-pass mode with default parameters. Only uniquely mapped reads were retained for downstream analyses. BAM files were sorted and indexed using samtools (v1.22.1). Gene read counts were quantified using featureCounts (v2.1.1) against GENCODE v49 gene annotations^72^. Only protein-coding genes were retained for downstream differential expression and pathway enrichment analyses. Differential gene expression analysis was performed in R (v4.4.1) using DESeq2 (v1.46.0)^73^. Genes with a log2(foldchange) ≥1 and adjusted P value *<* 0.05 (Benjamini– Hochberg correction) were considered significantly differentially expressed. Gene set enrichment and pathway analyses were performed using clusterProfiler (v4.16.0)^74,75^ against MSigDB Hallmark, GO BP and GO MF gene sets.

### Statistical analyses

All statistical analyses for *in vitro* and *in vivo* studies were performed using Prism 8.0 software (GraphPad). No data were excluded from the analyses. As sample sizes are similar to those previously reported^5,8^, no statistical methods were used to predetermine sample size, Specific statistical tests pertaining to each *in vitro* and *in vivo* experiment are described in the figure legends.

## Data availability

snRNA-seq data from the normal developing human cerebellum were obtained via correspondence from ref^4,28^ and are available through the Human Cell Atlas (https://www.covid19cellatlas.org/aldinger20), the UCSC Cell Browser (https://cbldev.cells.ucsc.edu) or from Database of Genotypes and Phenotypes (dbGaP) (accession number phs001908.v2.p1).

Curated independent single-cell MB datasets from previously published studies were obtained as follows: (1) Liu *et al* snRNA-seq data and scRNA-seq data were obtained via correspondence from Dr. Hailong Liu and are available through the China National GeneBank DataBase (CNGB) under accession numbers CNP0004439 and CNP0005220. (2) The full-length scRNA-seq data generated through Smart-seq2 from Hovestadt *et al*.^33^ was downloaded from (GEO) with the accession code GSE119926.

## Code availability

No custom code was developed for this study. Analyses were performed using open-source algorithms as described in the Methods. Additional details on the implementation and parameters are available from the corresponding authors upon reasonable request.

## Acknowledgements

We thank the ATC Gene Vector Core at Baylor College of Medicine for the preparation of lentiviral particles used in this study. We also thank M. “Sayeed” Sayeeduddin, Shahida Salar, and Zahida Sayeeduddin at the Pathology and Histology Core (HTAP) at Baylor College of Medicine for preparation of histological samples. T.E.W-O was supported by Texas Children’s Hospital, a Sam Day Foundation operating grant, a Dan L Duncan Comprehensive Cancer Center Mini-CiRCA grant, a Rally 20/20: Alumni Visionary Grant, and a Rally Foundation for Childhood Cancer Research/Kids Join the Fight Outside the Box Grant. T.E.W-O has also received generous support from Houston Tri Delta Philanthropies, Inc. O.S was supported by Enfants Cancers Santé (ECS) and the Société Française de Lutte contre les Cancers et les Leucémies de l’Enfant et de l’Adolescent (SFCE); with financial support from the 2021–2030 Cancer Control Strategy, on funds administered by Inserm; with support from the Institut national du cancer; and with support from the Ministère de l’Enseignement Supérieur et de la Recherche and the Agence Nationale de la Recherche. ELVN is a CPRIT Scholar in Cancer Research (RR200040). This project was partly supported by the Genomic and RNA Profiling Core at Baylor College of Medicine with funding from the NIH S10 grant (1S10OD036427), NIH NCI (P30CA125123), and CPRIT (RP250580) grants, and the Pathology and Histology Core (HTAP) with funding from P30 Cancer Center Support Grant (NCI-CA125123).

## Author contributions

R.S, X.Y, L.L (co-first author), and N.G designed, performed, and analyzed most of the experiments in the study. U.T performed intracerebellar transplants, and prepared xenograft samples for histology and sequencing. L.G.H and B.H contributed to the analyses of spliced variants by PCR and RBFOX2 OE experiments. L.L contributed to bulk and single nuclei RNA seq data analysis. M.B, C.A.LR, and A.C.S performed eCLIP and analyzed the datasets. I.B and O.A generated the RBFOX2 knockdowns for culture-adapted PDX7, culture-adapted PDX12 and ONS76 and collected the samples for further processing. E.L.V.N supervised the eCLIP studies, the associated bioinformatics analyses and figure contribution. R.S, X.Y, L.L (co-first author), O.S, J.Z and T.E.W-O prepared the figures and wrote the paper. O.S, J.Z. and T.E.W-O. jointly supervised the project.

