## Extended Data 1 to 9 and legends for "RBFOX2 restrains oncogenic cell plasticity in Group 3 medulloblastoma"

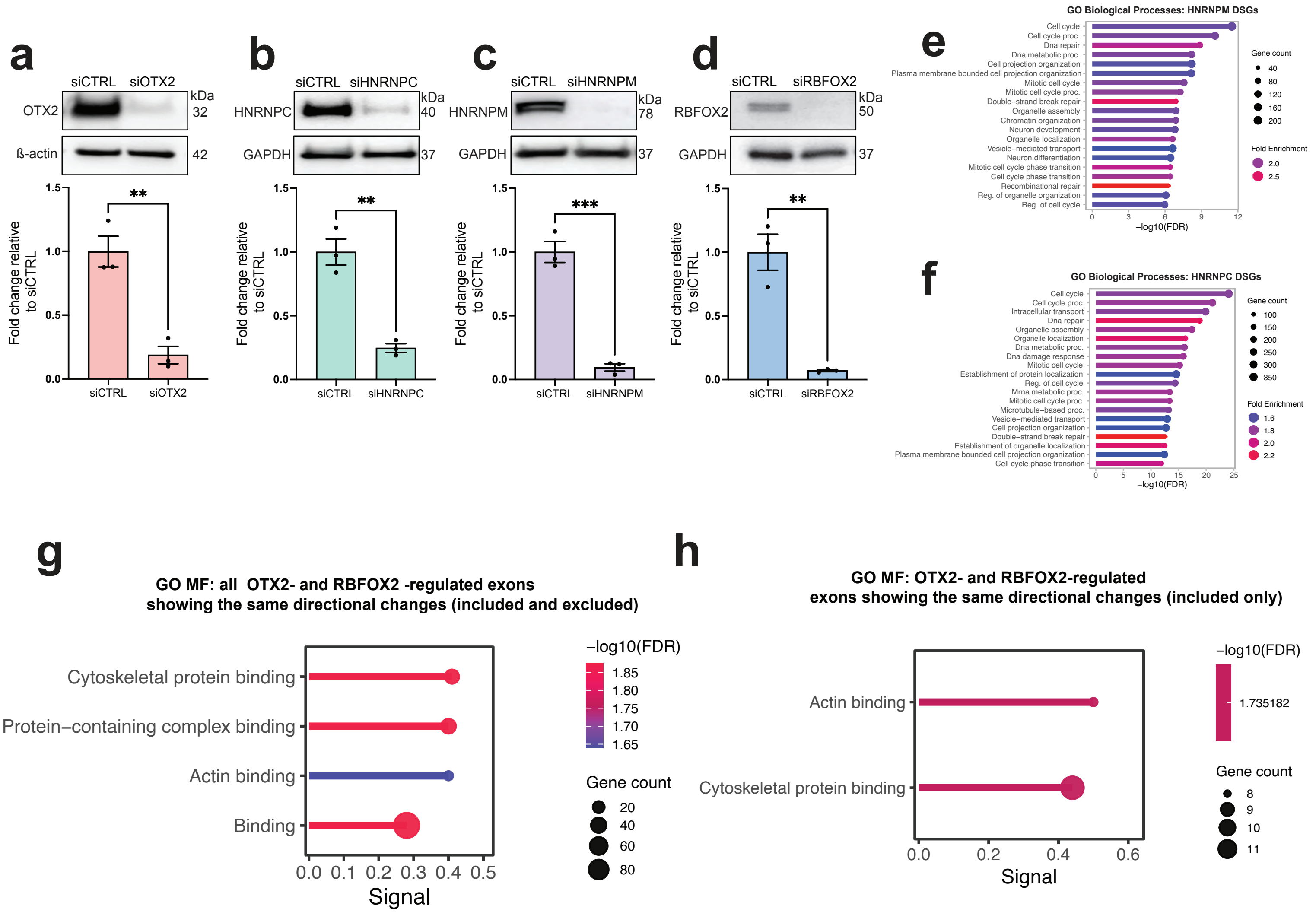

**Extended Data 1: RBFOX2 regulates divergent biological processes relative to HNRNPM and HNRNPC.**

a-d. Representative immunoblots (above) and quantification (below) of OTX2 (a), HNRNPC (b), HNRNPM (c) and RBFOX2 (d) silencing in HDMB03 tumorspheres. GAPDH or β-actin serve as loading controls. Error bars represent s.e.m (n=3 biological replicates per condition). Statistical significance was determined using an unpaired *t*-test. OTX2: **P=0.0043. HNRNPC: **P=0.0022. HNRNPM: P<0.001. RBFOX2: **P=0.0028.

e-f. GO Biological Processes (GO BP) for HNRNPM (e) and HNRNPC (f)-regulated differentially spliced genes (DSGs) demonstrating enrichment for cell cycle related processes.

g. GO Molecular Function (GO MF) enrichment of cytoskeletal dynamic properties for all OTX2- and RBFOX2-regulated alternatively spliced exons that are in the same direction (i.e exon included in both OTX2 and RBFOX2-regulated datasets or excluded in both datasets).

h. GO Molecular Function (GO MF) enrichment analysis depicting no significant enrichment for OTX2- and RBFOX2-regulated alternatively spliced exons in the same direction (included exons only).

### Extended Data 2

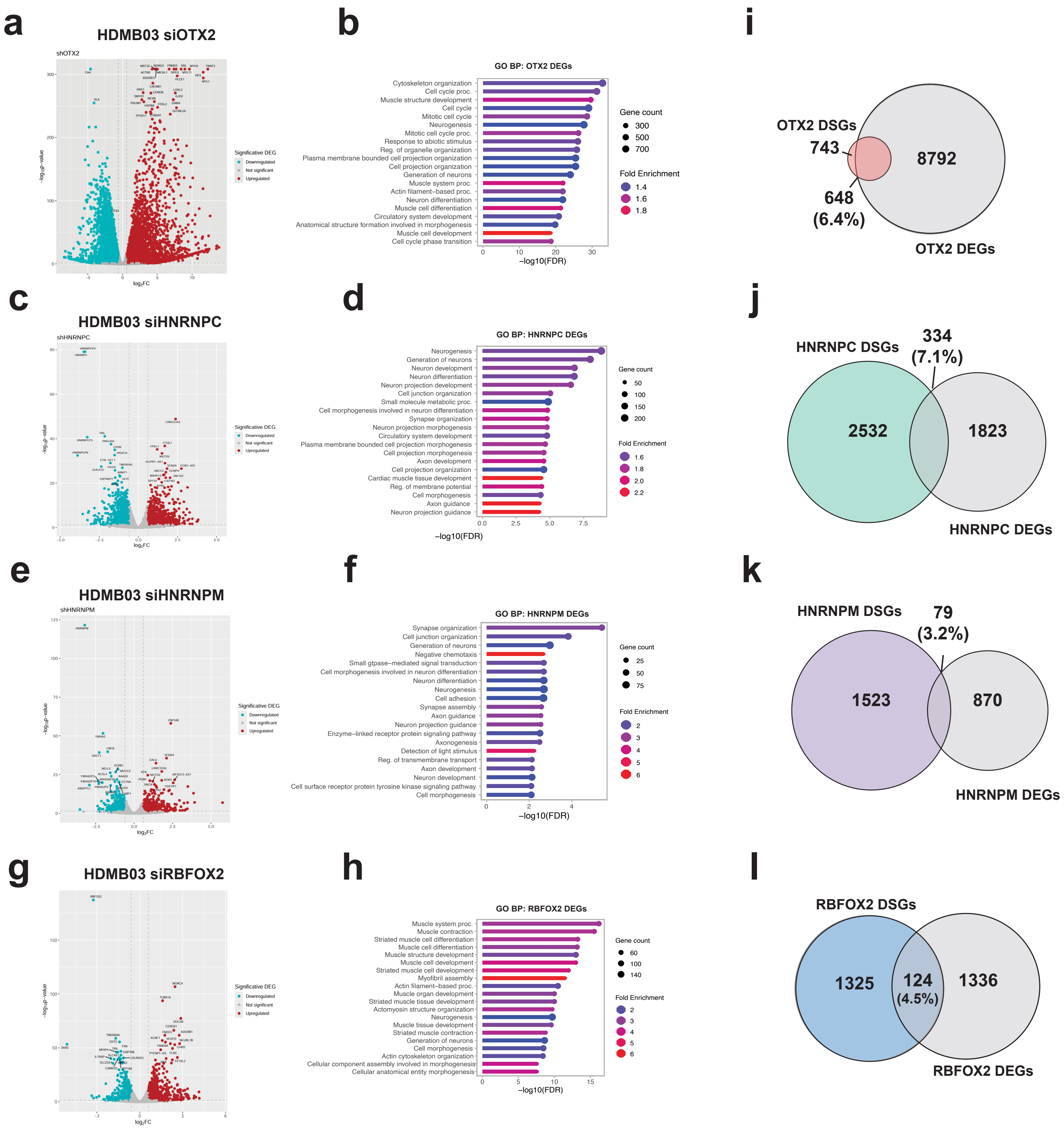

**Extended Data 2: Minimal overlap between OTX2 and LASR-regulated DEGs and DSGs in Group 3 MB tumorspheres.**

a. Volcano plot depicting significant and differentially expressed genes regulated by OTX2.

b. GO Biological Processes (GO BP) for OTX2-regulated differentially expressed genes (DEGs) demonstrating enrichment for cytoskeletal dynamics, neuronal differentiation and cell cycle.

c. Volcano plot depicting significant and differentially expressed genes regulated by HNRNPC.

d. GO Biological Processes (GO BP) for HNRNPC-regulated differentially expressed genes (DEGs).

e. Volcano plot depicting significant and differentially expressed genes regulated by HNRNPM.

f. GO Biological Processes (GO BP) for HNRNPM-regulated differentially expressed genes (DEGs).

g. Volcano plot depicting significant and differentially expressed genes regulated by RBFOX2.

h. GO Biological Processes (GO BP) for RBFOX2-regulated differentially expressed genes (DEGs).

i-l. Venn diagrams depicting percentage overlap between DSG and DEGs regulated by OTX2 (i), HNRNPC (j), HNRNPM (k) and RBFOX2 (l)

**Extended Data 3**

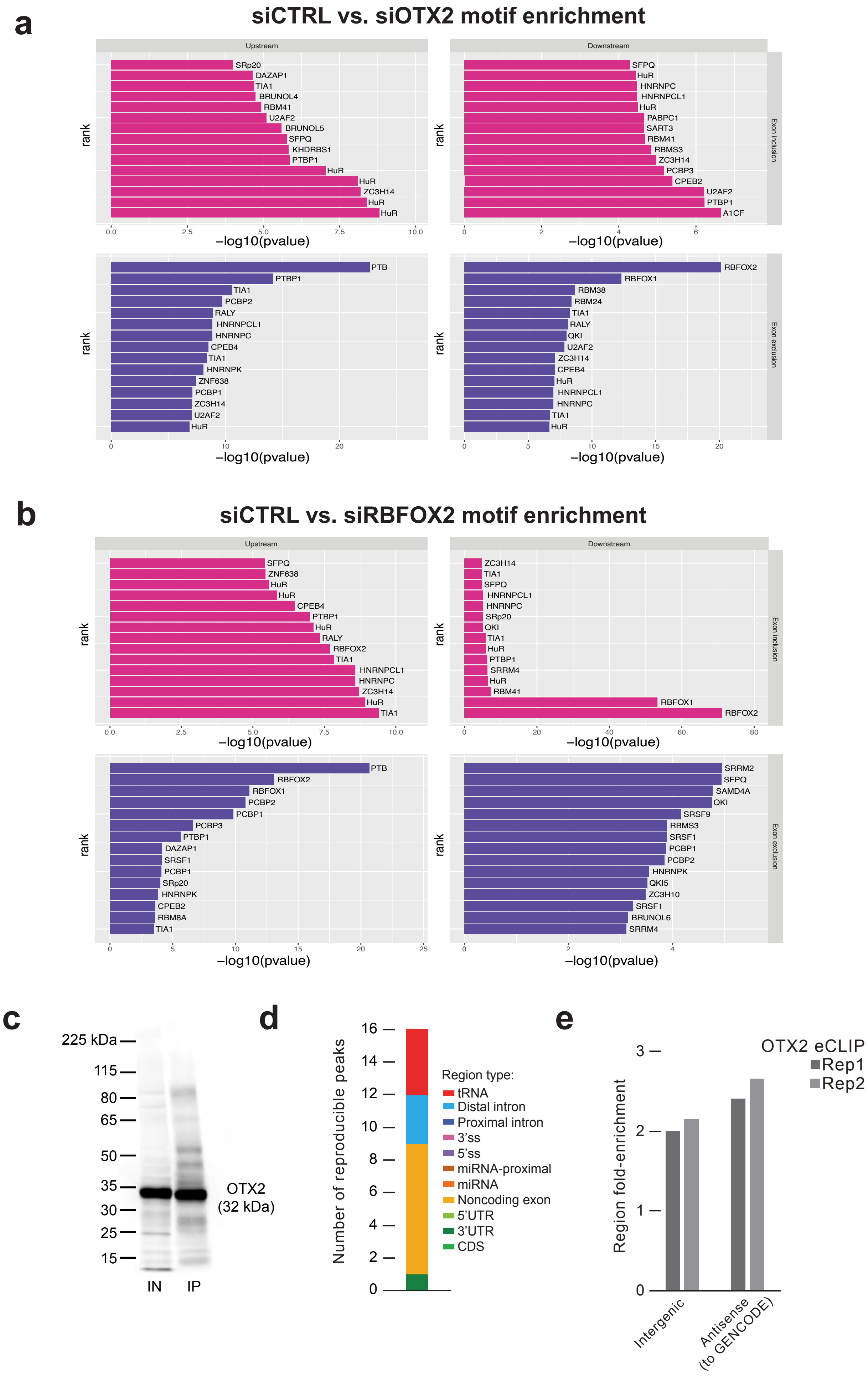

**Extended Data 3: OTX2 does not bind RNA to mediate alternative splicing events.**

a. RNA binding protein motif enrichment analysis in siCTRL vs siOTX2 conditions in HDMB03 Group 3 MB tumorspheres. RBFOX2 consensus motifs are significantly enriched downstream of exons that are excluded in siCTRL cells relative to siOTX2 cells. Top panels (pink) show motif enrichment upstream (left) or downstream (right) of included exons in siCTRL cells relative to siOTX2. Bottom panels (purple) show motif enrichment upstream (left) or downstream (right) of excluded exons in siCTRL cells relative to siOTX2.

b. RNA binding protein motif enrichment analysis in siCTRL vs siRBFOX2 conditions in HDMB03 Group 3 MB tumorspheres. RBFOX2 consensus motifs were significantly enriched downstream of exons that are included in siCTRL cells relative to siRBFOX2 cells. Top panels (pink) show motif enrichment upstream (left) or downstream (right) of included exons in siCTRL cells relative to siRBFOX2. Bottom panels (purple) show motif enrichment upstream (left) or downstream (right) of excluded exons in siCTRL cells relative to siRBFOX2.

c. Immunoblot depicting immunoprecipitation of OTX2 in HDBM03 tumorspheres. Input (IN) serves as a positive control for OTX2 immunoprecipitation (IP).

d. Stacked bar plot depicting minimal number of significant peaks identified across transcriptional loci in OTX2 IP Group 3 tumorspheres.

e. Plot depicting OTX2 binding to transcript intergenic regions. Rep = replicates.

**Extended Data 4**

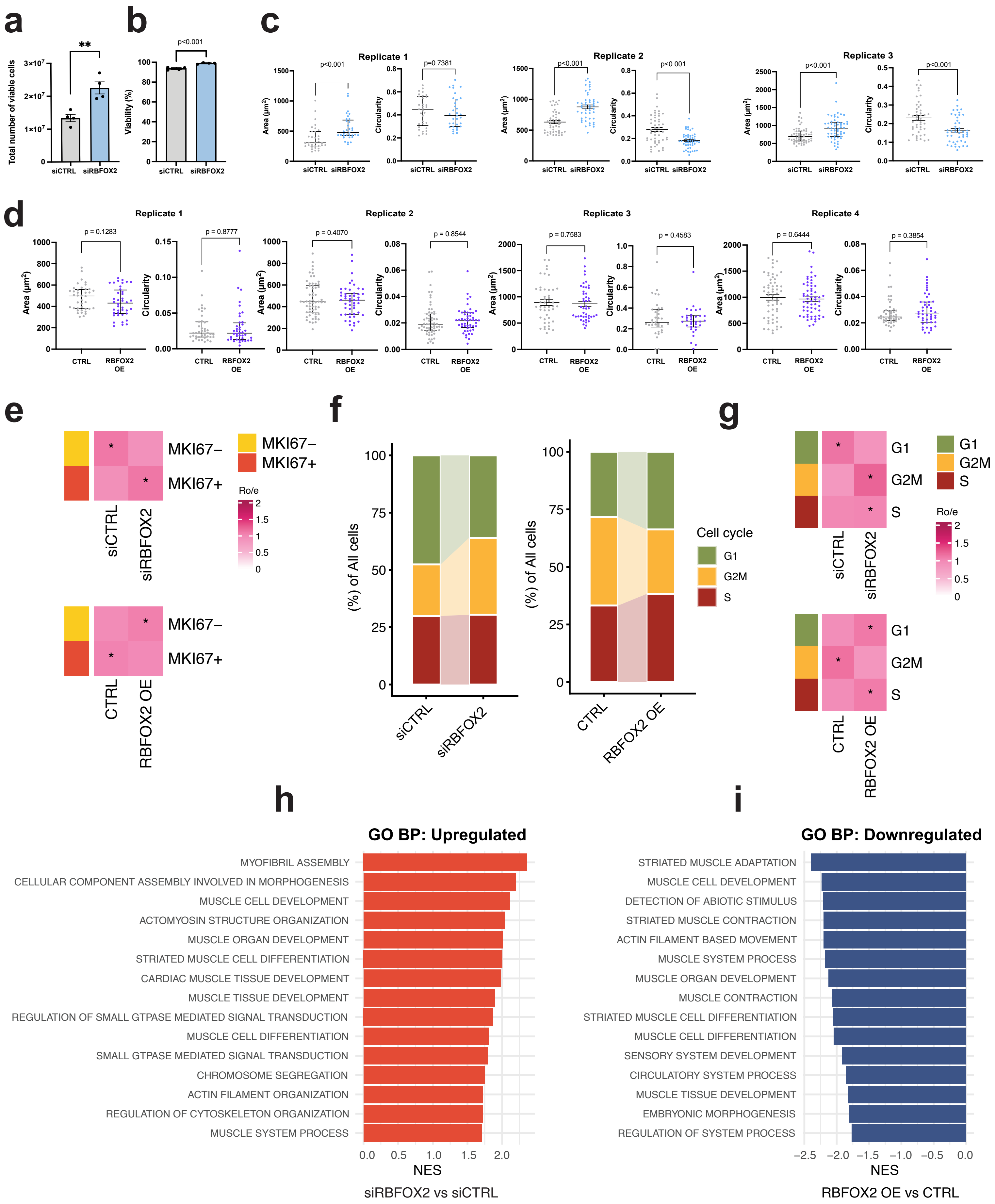

**Extended Data 4: RBFOX2 restrains Group 3 tumor cell plasticity *in vitro*.**

a-b. Quantification of cell number (a) and viability (b) in siRBFOX2 treated HDMB03 adherent cells compared to control. Error bars represent s.e.m (n=4 biological replicates). Statistical significance was determined using unpaired t-test. Cell number: **P=0.0053, Viability: P<0.001.

c. Quantification of cellular properties including area and circularity following phalloidin staining in siCTRL and siRBFOX2 treated HDMB03 cells. Results from three independent experiments. Error bars represent median with interquartile range. Statistical significance was determined using a Mann-Whitney test. Replicate 1: area; P <0.001, circularity; ns. Replicated 2: area; P <0.001, circularity; P<0.001. Replicate 3: area; P <0.001, circularity; P<0.001.

d. Quantification of cellular properties including area and circularity following phalloidin staining in CTRL and RBFOX2 OE treated HDMB03 cells. Results from four independent experiments. Error bars represent median with interquartile range. Statistical significance was determined using a Mann-Whitney test. Replicate 1: area; P = 0.1283, circularity; p=0.8777. Replicate 2: area; P =0.4070, circularity; P=0.0.8544. Replicate 3: area; P=0.7583, circularity; P=0.4583. Replicate 4: area; P=0.6444, circularity; P=0.3854.

e. Group prevalence of proliferation cell states measured by Ro/e between siCTRL and siRBFOX2 (top), and between CTRL and RBFOX2 OE (bottom).

f. Composition of cell cycle phases between siCTRL and siRBFOX2 (left), and between CTRL and RBFOX2 OE (right).

g. Group prevalence of cell cycle phases measured by Ro/e between siCTRL and siRBFOX2 (top), and between CTRL and RBFOX2 OE (bottom).

h. Top 15 significantly enriched upregulated GO Biological Process (GO BP) pathways in siRBFOX2 versus siCTRL cells.

i. Top 15 significantly enriched downregulated GO Biological Processes (GO BP) pathways in RBFOX2 OE versus CTRL cells.

### Extended Data 5

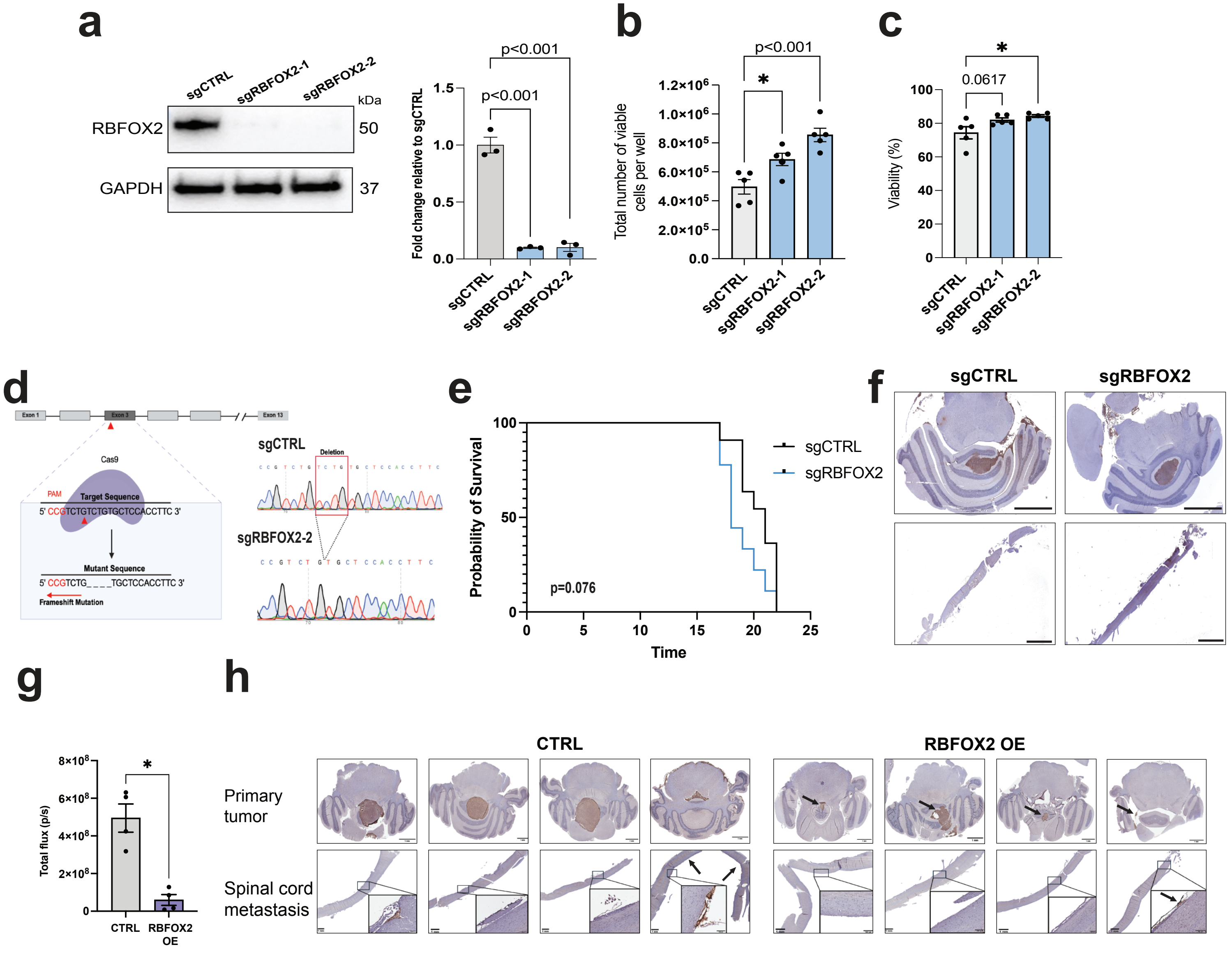

**Extended Data 5: RBFOX2 overexpression inhibits tumor progression *in vivo*.**

a. Immunoblots (left) and quantification (right) depicting RBFOX2 loss following CRISPR knockout using 2 sgRNAs targeting *RBFOX2* exon 2 (sgRBFOX2-1) or exon 3 (sgRBFOX2-2) in HDMB03 cells. GAPDH serves as a loading control. Statistical significance determined using one-way ANOVA followed by a Dunnett's test for multiple comparisons. P<0.001 for both sgRNAs.

b-c. Quantification of cell number (b) and viability (c) in sgRBFOX2-1 and sgRBFOX2-2 knockout HDMB03 cells relative to sgCTRL. Error bars represent s.e.m. (n=5 biological replicates). Statistical significance determined using one-way ANOVA followed by a Dunnett's test for multiple comparisons. Cell number: *P=0.0249 for sgRBFOX2-1 and ***P=0.0003 for sgRBFOX2-2. Viability: P=0.0617 for sgRBFOX2-1 and *P=0.0163 for sgRBFOX2-2.

d. Schematic depicting methodology used to generate CRISPR-guided RBFOX2 knockout (sgRBFOX2) and non-targeting control (sgCTRL) HDMB03 cells that were injected into the cerebellum of NSG mice. A single guide RNA (sgRNA) targeting exon 3 of the RBFOX2 gene resulted in a 4 base pair deletion and subsequent frameshift mutation to knockout RBFOX2 expression.

e. Kaplan–Meier curves following transplantation of NSG mice with sgCTRL HDMB03 cells (n=11) and sgRBFOX2 HDMB03 cells (n=10). Statistical significance was determined by a log-rank test. P=0.076.

f. Representative immunohistochemistry (IHC) staining for mitochondria to detect human cells from animals injected with $1.5x10^4$ sgCTRL or sgRBFOX2 cells at Day 15 post injection. Scale bars: 4 mm.

g. Quantification of tumor volume in animals injected with $1.5x10^3$ CTRL and RBFOX2-OE HDMB03 cells, calculated by BLI total flux (p/s). Statistical significance was determined using a Mann-Whitney test. n=4 per group, *P=0.0286.

h. Immunohistochemistry (IHC) staining for mitochondria to detect human cells from animals injected with $1.5x10^3$ CTRL or RBFOX2 OE cells demonstrating the difference in primary tumor volume at Day 15 post injection. Scale bars: 1 mm.

#### Extended Data 6

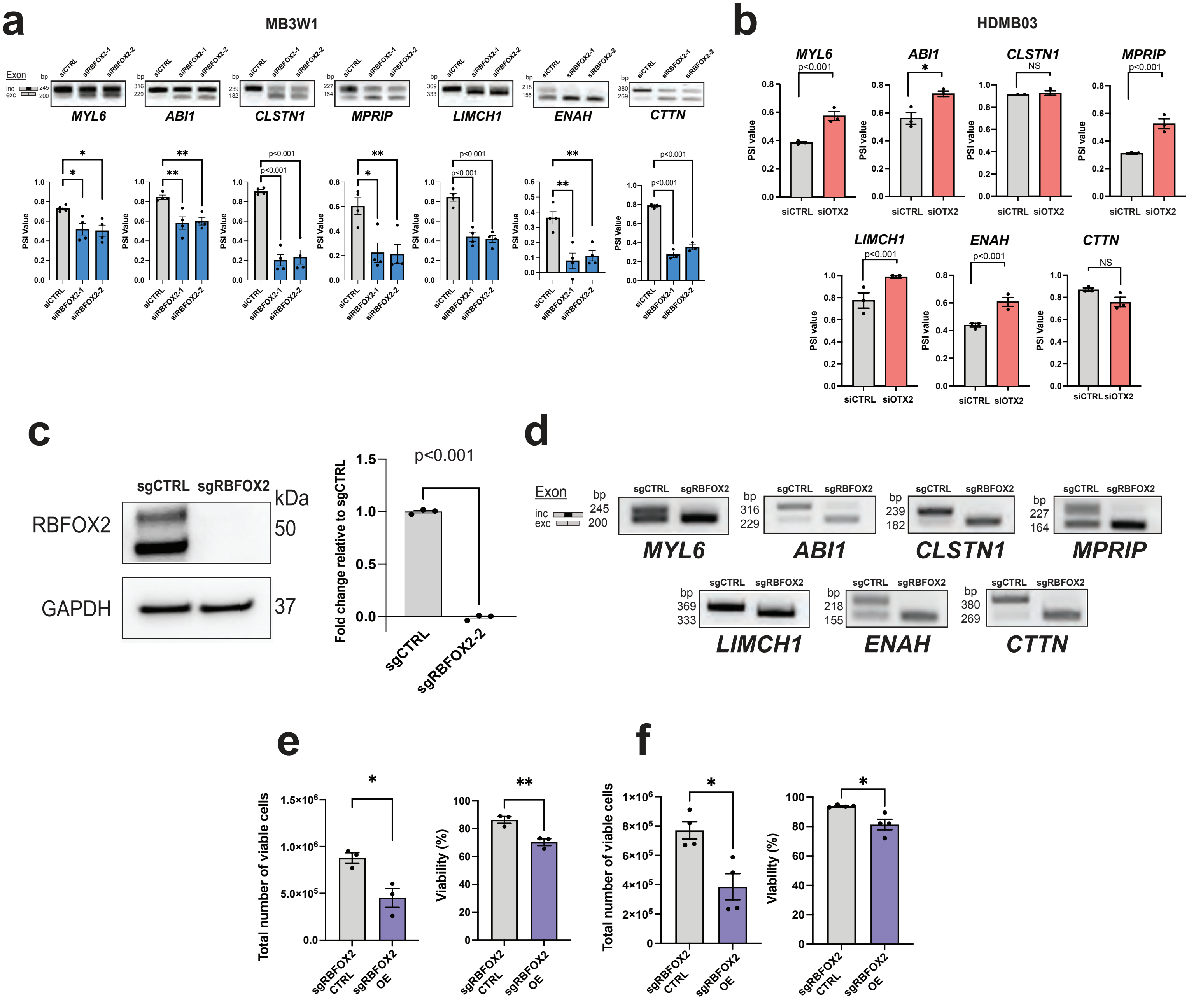

**Extended Data 6: Cytoskeletal-associated gene splicing is directly dependent on RBFOX2, including genes associated with Rac1-WAVE-Arp2/3 signaling.**

a. RT-PCR validation (upper) and associated quantification (lower) of alternatively spliced exon levels for select cytoskeletal remodeling genes in RBFOX2 silenced MB3W1 tumorspheres. Evaluation carried out with 2 independent RBFOX2 siRNAs. Error bars represent s.e.m. n=4 biological replicates for each gene. Statistical significance determined using an unpaired t-test. *MYL6*: siRBFOX2-1; *P=0.0235, siRBFOX2-2; *P=0.0166. *ABI1*: siRBFOX2-1; **P=0.004, siRBFOX2-2; **P=0.006. *CLSTN1*: siRBFOX2-1; P<0.001, siRBFOX2-2; P<0.001. *MPRIP*: siRBFOX2-1; *P=0.0112, siRBFOX2-2; **P=0.0095. *LIMCH1*: siRBFOX2-1; P<0.001, siRBFOX2-2; P<0.001. *ENAH*: siRBFOX2-1; **P=0.0019, siRBFOX2-2; **P=0.0041. *CTTN*: siRBFOX2-1; P<0.001, siRBFOX2-2; P<0.001.

b. Quantification of splicing alterations of select cytoskeletal remodeling genes from RNA-sequencing following OTX2 silencing in HDMB03 tumorspheres. Error bars represent s.e.m. (n=3 biological replicates). Statistical significance determined using unpaired t-test. *MYL6*; P<0.001, *ABI1*; *P=0.046, *CLSTN1*; NS, *MPRIP*; P<0.001, *LIMCH1*; P<0.001, *ENAH*; P<0.001, *CTTN*; NS.

c. Representative immunoblot (left) and quantification (right) confirming sustained loss of RBFOX2 expression in CRISPR-guided RBFOX2 knockout HDMB03 cells following clonal expansion to remove cells with residual RBFOX2. GAPDH serves as a loading control. Error bars represent s.e.m. (n=3). Statistical significance determined using unpaired t-test. P<0.001.

d. RT-PCR demonstrating sustained exon skipping events in RBFOX2 CRISPR knockout (sgRBFOX2) HDMB03 cells relative to controls (sgCTRL) following clonal expansion.

e. Quantification of cell number (left) and viability (right) following RBFOX2 overexpression in the clonal sgRBFOX2 knockout (sgRBFOX2 OE) tumorspheres compared to sgRBFOX2 knockout control cells (sgRBFOX2 CTRL). Error bars represent s.e.m. n=3 biological replicates. Statistical significance determined using an unpaired t-test. Cell number; *P=0.0206, viability; **P=0.0095

f. Quantification of cell number (left) and viability (right) following RBFOX2 overexpression in the clonal sgRBFOX2 knockout (sgRBFOX2 OE) adherent cells compared to sgRBFOX2 knockout control cells (sgRBFOX2 CTRL). Error bars represent s.e.m. n=4 biological replicates. Statistical significance was determined using an unpaired t-test. Cell number; *P=0.0113, viability; *P=0.0135.

#### Extended Data 7

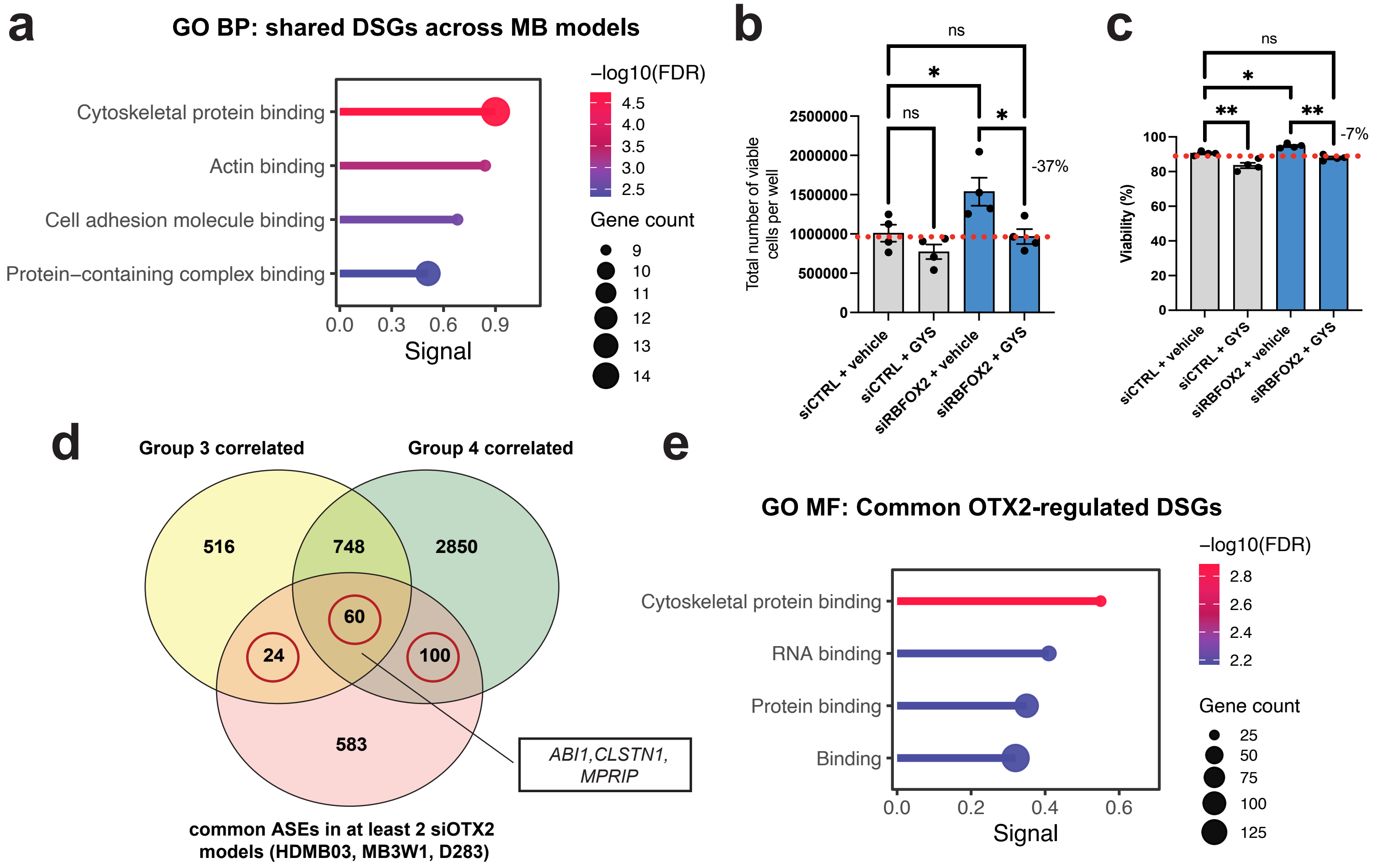

**Extended Data 7: RBFOX2-mediated AS converges on Rac1-dependent cytoskeletal machinery.**

a. Enrichment of GO Biological Processes (GO BP) for the 42 shared ASEs conserved across MB models, demonstrating enrichment of pathways associated with cytoskeletal dynamics.

b-c. Quantification of cell number (b) and viability (c) in siCTRL and siRBFOX2 cells following 20 μM Rac1 inhibitor treatment. Error bars: s.e.m. n =4 biological replicates. Note the 37% drop in cell number for treated siRBFOX2 cells is not due to cell death, as there was a minimal decrease in viability. siCTRL + vehicle vs. siRBFOX2 + vehicle cell number: *P=0.0468, siRBFOX2 + vehicle vs. siRBFOX2 + GYS cell number: *P=0.0307. siCTRL + vehicle vs. siRBFOX2 + vehicle viability: *P=0.0455, siCTRL + vehicle vs. siCTRL + GYS viability: **P=0.0016. siRBFOX2 + vehicle vs. siRBFOX2 + GYS viability: **P=0.0018.

d. Venn diagram showing overlap between significant OTX2-regulated ASEs in Group 3 MB models (HDMB03, MB3W1 and D283) and OTX2-correlated exons in Group 3 MB and Group 4 MB patient samples, which includes core cytoskeletal proteins *ABI1*, *CLSTN1*, and *MPRIP*.

e. GO Molecular Function (GO MF) enrichment for shared significantly OTX2-regulated ASEs in Group 3 MB models and OTX2-correlated exons in Group 3 MB and Group 4 MB patient samples.

#### Extended Data 8

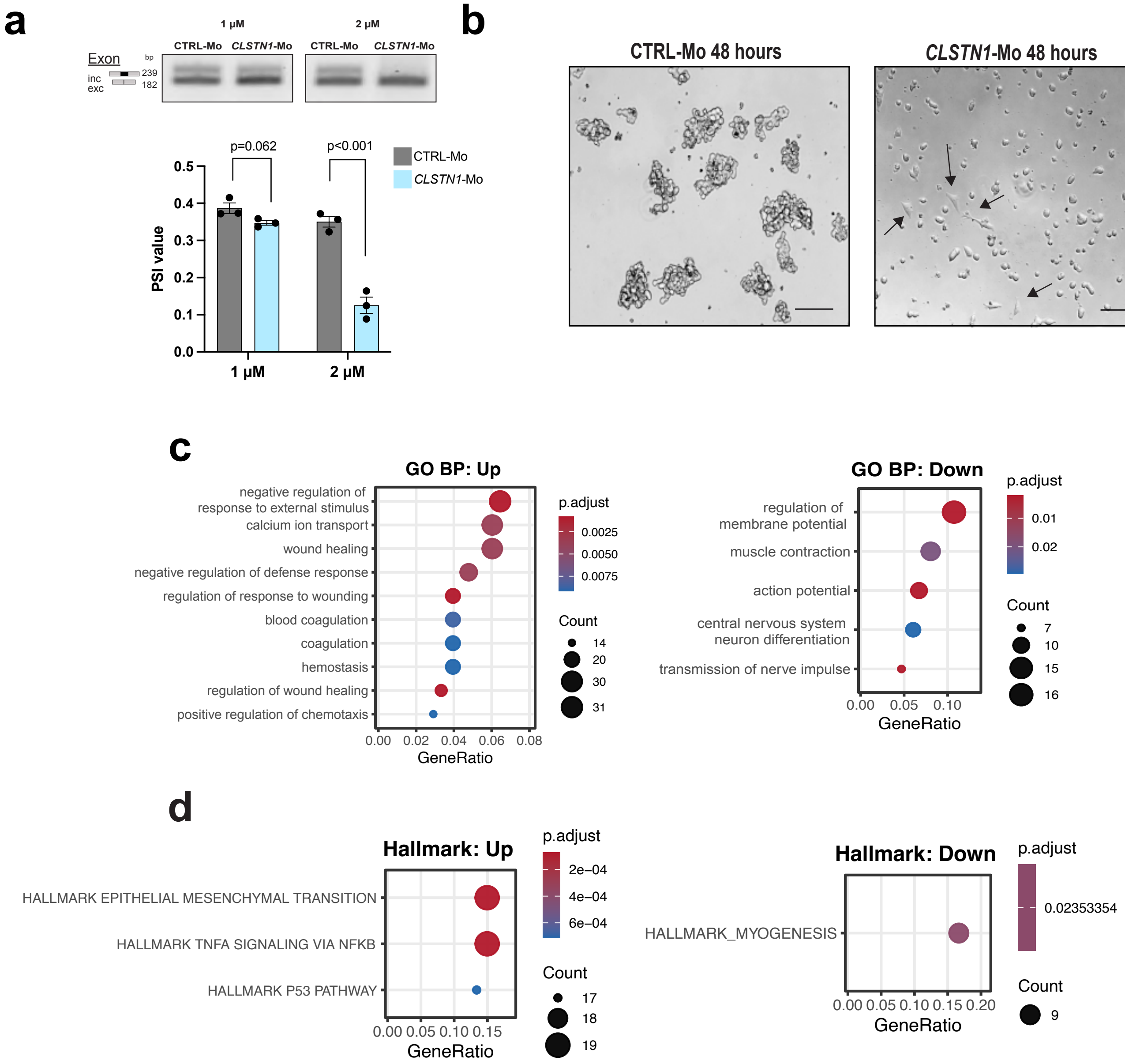

**Extended Data 8: Morpholino-induced exon skipping leads to morphological and transcriptional changes associated with RBFOX2 loss.**

a. RT-PCR validation (upper) and associated quantification (lower) of *CLSTN1* exon exclusion following 48 hours of 1 μM and 2 μM morpholino (Mo) treatment in HDMB03 cells. Error bars represent s.e.m. (n=3 biological replicates for each condition). Statistical significance determined using unpaired t-tests. 1 μM; $P=0.062$, 2 μM; $P<0.001$. PSI = percent spliced in.

b. Representative images of 2 μM Mo-treated cells at 48 hours depicting an increase in cell-substrate adhesion. Scale bar: 150 μm.

c. Top 10 enriched GO Biological Process (GO BP) pathways from significantly upregulated (left) and downregulated (right) genes in *CLSTN1*-Mo versus CTRL-Mo samples. Dot size indicates the number of mapped genes, and color indicates the adjusted P value.

d. Significantly enriched Hallmark pathways from significantly upregulated (left) and downregulated (right) genes in *CLSTN1*-Mo versus CTRL-Mo samples. Dot size indicates the number of mapped genes, and color indicates the adjusted P value.

### Extended Data 9

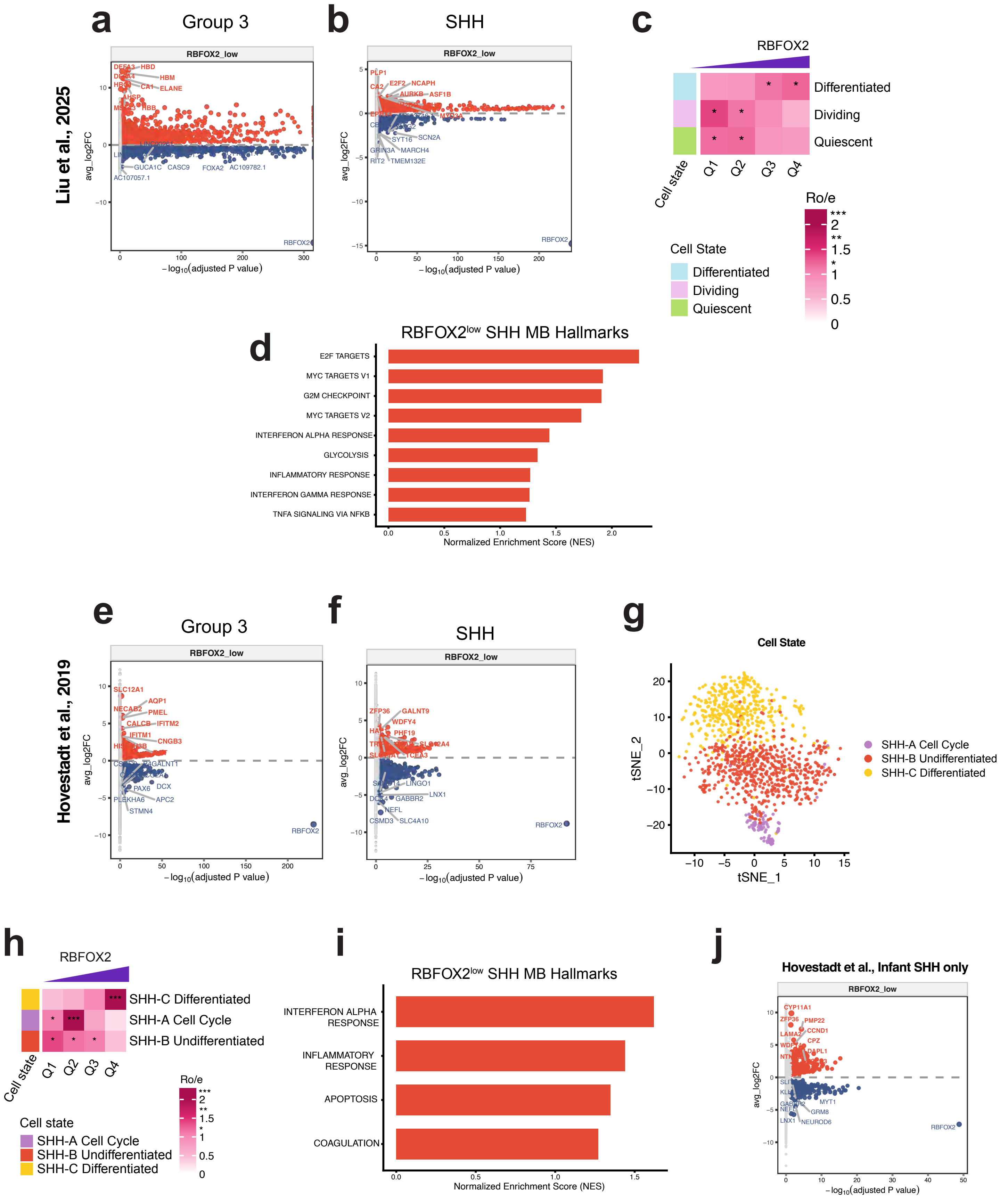

**Extended Data 9: RBFOX2-mediated cell state extends to SHH MB patient tumor cells.**

a-b. Volcano plots of differentially expressed genes comparing RBFOX2$^{low}$ versus RBFOX2$^{high}$ tumor cells in Group 3 MB (a) and SHH MB (b) from the Liu et al. cohort. Upregulated genes are shown in red and downregulated genes in blue.

c. Cell state composition across RBFOX2 expression quartiles (Q1–Q4, lowest to highest) in SHH MB dataset from the Liu et al. cohort.

d. Hallmark gene set enrichment analysis (GSEA) comparing RBFOX2$^{low}$ and RBFOX2$^{high}$ tumor cells in the SHH MB dataset from the Liu et al. cohort.

e-f. Volcano plots of differentially expressed genes comparing RBFOX2$^{low}$ versus RBFOX2$^{high}$ tumor cells in Group 3 MB (e) and SHH MB (f) from the Hovestadt et al. cohort. Upregulated genes are shown in red and downregulated genes in blue.

g. UMAPs shows the single cell profiles of SHH MB dataset from the Hovestadt et al. cohort, with cells colored by cell states.

h. Cell state composition across RBFOX2 expression quartiles (Q1–Q4, lowest to highest) in the SHH MB dataset from the Hovestadt et al. cohort.

i. Hallmark gene set enrichment analysis (GSEA) comparing RBFOX2$^{low}$ and RBFOX2$^{high}$ tumor cells in the SHH MB dataset from the Hovestadt et al. cohort.

j. Volcano plots of differentially expressed genes comparing RBFOX2$^{low}$ versus RBFOX2$^{high}$ tumor cells in SHH MB infant patients from the Hovestadt et al. cohort. Upregulated genes are shown in red and downregulated genes in blue.
